# Artificial intelligence-based gas chromatography-olfactometry for sensory evaluation of key compounds in food ingredients

**DOI:** 10.1101/2022.04.20.488977

**Authors:** Liang Shang, Chuanjun Liu, Fengzhen Tang, Bin Chen, Lianqing Liu, Kenshi Hayashi

## Abstract

The use of an artificial intelligence (AI)-based prediction model of the structure-odor relationship (SOR) has shown great potential in the replacement of human panelists in gas chromatography-olfactometry (GCO). However, the Al-based GCO encounters issues such as poor accuracy, generalization, and practicality, owning to the insufficient feature extraction of odorant molecular structure. The purpose of this study is to the prediction of odor perception categories based on the odorant structure feature extraction by diverse deep neural networks, including molecular graphic convolution neural network (MG-CNN), molecular graph transformer neural network, and atom interaction neural networks. The results of the performance comparison of different feature extractors demonstrate that the MG-CNN model produces the highest accuracy and thus may be most suitable for the SOR prediction. It is hoped that the proposed method can be applied in practice as an auxiliary tool of GCO for the sensory evaluation of key compounds in food ingredients.

**Highlights:**

- Different deep neural networks were used to predict categorized odor descriptors.
- End-to-end-based representation learning was performed for molecular feature extraction.
- Molecular graphs with pre-training of convolution neural networks was most accurate.

**Graphical Abstract:** 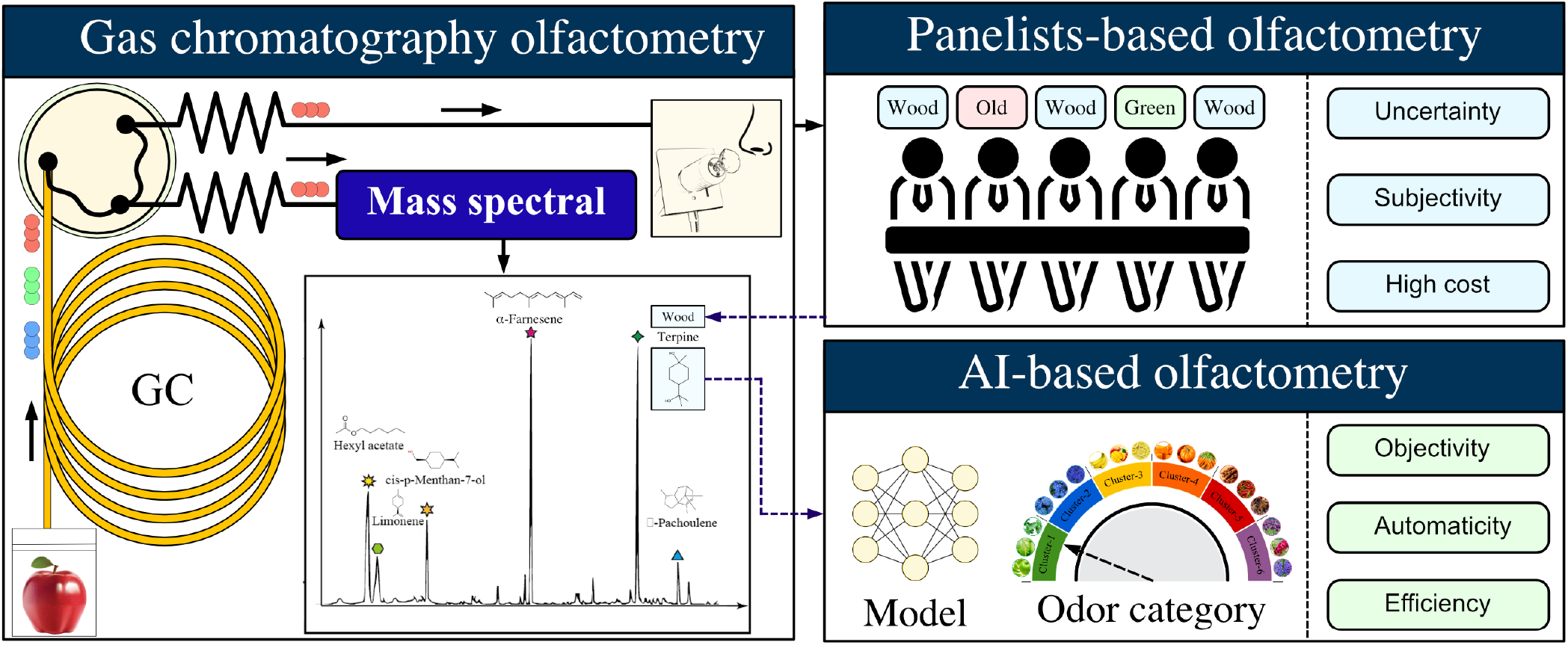

## 1. Introduction

Characterization of the key aroma-active compounds from food is an important topic in food science. To obtain the sensory information of odor, gas chromatography-olfactometry (GCO) has been widely applied as a powerful odor analytic strategy in various research areas, especially in food science (Maurya et al., 2021). Recently, GCO combined with headspace solid-phase microextraction (HS-SPME) has been employed to determine the key aroma compounds of tea (Ma et al., 2022), coffee (Caporaso et al., 2022), pork (Wu et al., 2022), Baijiu (Huang et al., 2022), spices (Ni et al., 2022), and celeries (Turner et al., 2021) et al.

Although GCO can be used to attain accurate sensory and chemical characteristics of odorants, this approach is expensive and time-consuming, which can limit its application. The main expense associated with GCO is the cost of hiring and training panelists to characterize the sensory qualities of odorants using their sense of smell (Acree, 1997). In addition, the sensory assessment of human panelists is personally dependent, which inevitably leads to subjectivity and inconsistency into the evaluation results. In order to overcome the above problems, some auxiliary business software modules, such as GC/MS Off-Flavor Analyzer (Shimadzu Co. Ltd.), AromaOffice (GERSTEL K.K.), and AroChemBase (Alpha MOS Co. Ltd.), have been widely applied as supplements to GCO analysis. The number of odorants with odor descriptors (around 2000 compounds for AroChemBase, 600 compounds for GC/MS Off-Flavor Analyzer) that are collected in their aroma compound databases. However, there are around 44,000 odorants in the world, and most of them are unknown to us. It indicated that existing systems may not meet the harsh requirements of GCO in food aroma analysis. Recently, the use of artificial intelligence (AI) to support analytical purposes has been a potential technology for overcoming the drawbacks of traditional analytical methods. Therefore, AI-based auxiliary for human assessors would reduce the above problems.

Research on the response patterns of neurons in the olfactory bulbs (OB) has illuminated the mechanisms underlying biological olfaction. However, many questions remain, such as why these molecules smell different from one another and why we link smell feelings with certain semantic descriptors, known as odor descriptors (ODs) (Barwich, 2020). These questions may be answered by examining structure odor relationships (SORs), the development of which presents a difficult and interesting challenge. In recent years, many types of research have been conducted to predict odor perception of odorants using various parameters, such as electronic or physicochemical characteristics (Chacko et al., 2020), mass spectrometry (MS) (Nozaki & Nakamoto, 2018), and social network interactions (Snitz et al., 2019). Additionally, novel methods, such as odor-based social networks (Kumar et al., 2015; Snitz et al., 2013; Liu et al., 2019), machine learning (ML) (Loetsch et al., 2019; Choi et al., 2022), deep neural network (DNN) models (Debnath & Nakamoto, 2020), and semantic-based approaches (Gutiérrez et al., 2018) have been developed to calibrate models that express the relationships between odorants and ODs. These studies have demonstrated the possibility to use data-driven approaches to solve the SOR problems. In line with this trend, we have proposed a concept of ML-based GC/MS olfactometry in which the sensory evaluation of panelists is expected to be replaced by machine learning prediction models (Shang et al., 2017).

In the ML-based GCO system, the molecular information obtained by MS analysis of the individual GC peaks is transfered into physicochemical parameters by molecular calculation software (DRAGON). By building models appropriately, the ODs of the odorants can be predicted with high accuracy from their physicochemical parameters. The ML-based GCO is a concept-of-proof study and there are many problems to be solved before its practical application. For example, in terms of the sample size, only limited ODs with a high frequency of occurrence are targeted in our models. In reality, it has been reported that over 500 ODs are used for odor evaluation (Dravnieks, 1982). Therefore, many ODs, especially for those rare ODs are not considered by the models. In order to solve this issue, we have recently proposed a categorization approach based on the semantic analysis of ODs, which make the model can cover the prediction of several hundreds of ODs (Shang et al., 2021). Another remaining problem is that the model prediction is based on physicochemical parameters of odorants. The recent development of computational chemistry and ML makes it possible to obtain various molecular structure information (Tsubaki & Mizoguchi, 2019). Unlike traditional methods, DNNs can directly learn latent presentations from molecular structures, such as atom types and their positions, via back-propagation. Moreover, an end-to-end strategy has been proposed as an effective nonlinear modeling method for learning molecular presentation in many fields, such as quantum chemical properties prediction, and compound-protein interaction identification (Sharma et al., 2021). However, since the detailed binding mechanism between the odorants and olfactory receptors is still not fully understood, which molecular features play the most important role in olfactory perception has not been cleared.

The purpose of this study is dedicated to molecular feature mining by diverse DNNs for the prediction of odor perception categories. We focus on using multi-type of feature extractors to understand the relationship between the odorant structure features and odor perception categories and thus construct a prediction model with much higher accuracy. This will make the ML-GCO system more generalizable and therefore more practical. A schematic of the data processing and modeling procedures is illustrated in Fig. 1. We used molecular structures, including 2D molecular images and atom spatial locations, to predict odor sensory categories. For minding useful information from molecular structures, some molecular structural feature extraction methods were applied and their results were compared and discussed. Specifically, we used a molecular graphic convolution neural network (MG-CNN) and molecular graph transformer neural network (MGTNN) to extract features from representations of molecular structure, such as molecular structure images and topology graphs. Moreover, we used atom interaction neural networks (AINNs) to generate features from the spatial structures of atoms. To overcome limitations related to insufficient samples, we also considered pre-trained models, such as pre-trained CNNs and MGTNNs. We compared the feasibility of SOR prediction via molecular fingerprints, molecular parameters, and molecular feature extraction based on MG-CNN, MGTNN, and AINN. We achieved the highest performance by applying molecular features extracted via a pre-trained MG-CNN combined with a multi-label learning model (area under the ROC curve: 0.877±0.028 and F1-score: 0.726±0.028). It indicated that molecularly image features would provide more sensory information than others. Thus, the proposed multi-label odor sensory category identification model is a feasible option for developing AI-based GCO. Although it would be still a challenge to replace human assessors with ML models perfectly, the proposed method could provide references for assessors to increase the efficiency of food aroma analysis.

**Figure 1:**
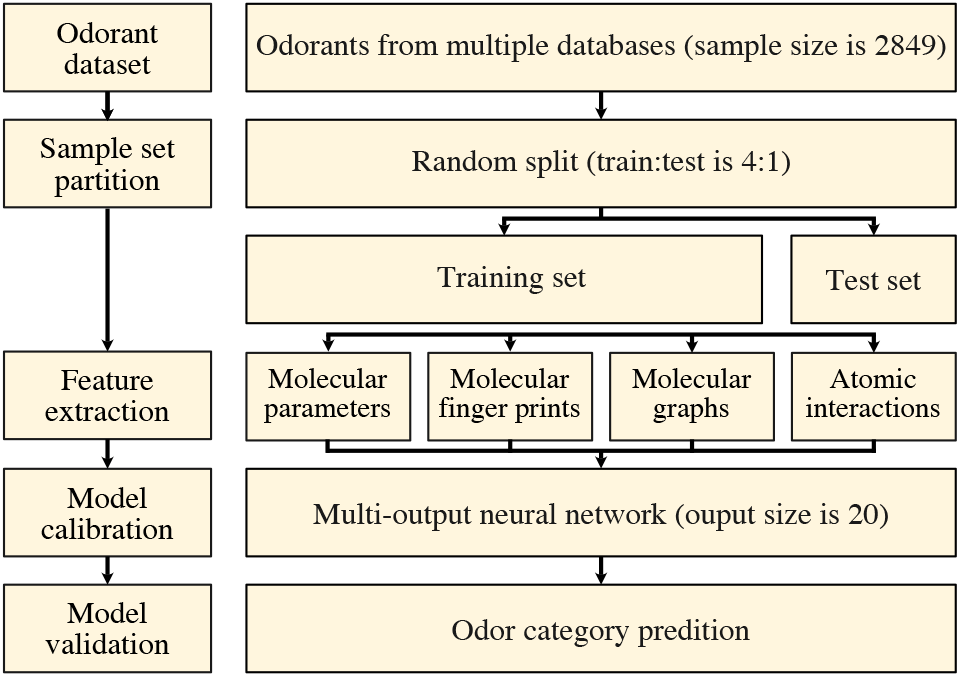
Data processing for calibration and validation of an odor category prediction model.

## 2. Materials and methods

### 2.1. Data collection and preparation

To create an odor category prediction model, we collected an extensive dataset containing 2849 odorants. We used information from publicly available databases such as the Odor Map database (Johnson et al., 2009), Flavors and Fragrances database (Sigma-Aldrich) (Sigma-Aldrich, 2016), and the Good Scents database (Arn et al., 1998). As illustrated in Fig. 1, we collected and prepared 5 types of molecular descriptors, including molecular parameters, molecular fingerprints, molecular graphs, simplified molecular-input line-entry system (SMILES), and atom coordinates. Specifically, we used SMILES and RDKit Software (ver. 2021.03.1) to collect normalized molecular parameters (1826), molecular fingerprints (binary, 2048 bits), and 2D molecular graphs (RGB, 300×300 pixels). We used RDKit to obtain the 2D and 3D atom coordinates of the odorants, which we used to train the model regarding the atomic interactions between odorant structures and their odor sensory categories. In total, 256 ODs were clustered into 20 categories using a co-occurrence Bayesian embedding method. More detailed information regarding the cleaning and categorization of ODs can be found in Table S1 (Shang et al., 2021). Data were processed and analyzed using Python (ver. 3.9.0) and R (ver. 4.1.1). Because of the diverse molecular representations of odorants, we need to employ various feature extraction technologies to obtain embeddings for model calibration.

### 2.2. Model calibration

The calibration and validation process for the odor category model is shown in Fig. 1. First, all samples were divided via random splitting into training and test sets with a 4:1 ratio, and the reported results are averaged over 50 repetitions. Afterward, we employed molecular parameters, molecular fingerprints, molecular graphic features, molecular graph transformers, and atomic interaction embedding to extract the molecular features of the odorants (Fig. 2). We used a DNN approach to develop a multi-label odor category learning model (the output size was 20) based on the afore-mentioned molecular information. The cost function ℒ(Θ) of the DNN multi-label classifier was calculated by summing the binary cross-entropy of each class, which was defined as follows:

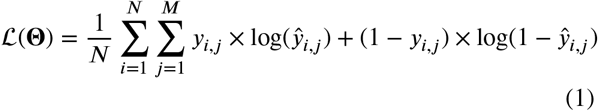

where Θ indicates the parameter set of the model, *N* indicates the sample number in the training set, and *M* indicates the number of odor categories, set as 20 in the present study. *y*_*i,j*_ and ŷ_*i,j*_ are the ground truth and prediction label for category *j* of sample *i*, respectively. By minimizing the cost function based on the stochastic steepest gradient descent algorithm, parameters from the DNN could be learned and updated. For each DNN method, the number of hidden layers and nodes was selected from 1 to 8 layers and {16, 32, 64, 128, 256, 512, 1024, 2048} nodes, respectively. Network parameters, including the dropout ratio, learning rate, and training epoch, were set as 0.1, 1 × 10^−4^, and 200, respectively. The optimal models were determined according to their areas under the ROC curve (AUC) and F1-scores based on precision and recall, simultaneously. Both qualitative and quantitative data analyses were performed.

**Figure 2:**
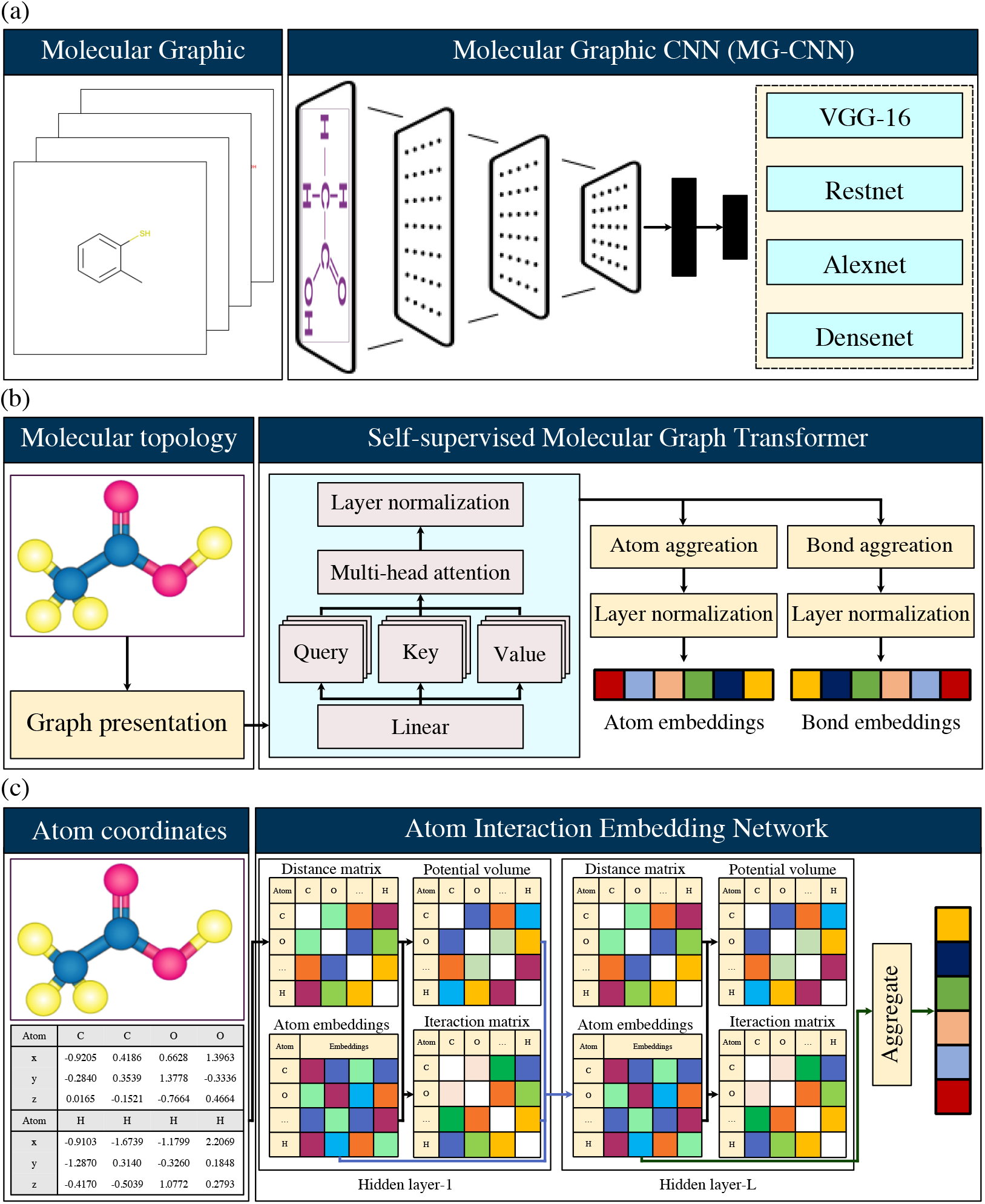
Overview of the design of molecular feature extractors, including (**a**) a molecular graphic convolution neural network (MG-CNN), (**b**) molecular graph transformer neural network (MGT-NN), and (**c**) atom interaction neural network (AINN).

Because molecular graphs, molecular SMILES sequences, and atom coordinates are not tabular data, they cannot be used as classifier inputs directly. Therefore, diverse feature extraction methods were first employed to convert those unstructured data to tabular features. Specifically, pre-trained CNNs, sequence transformer, and atomic interaction embedding were utilized for molecular graphs, SMILES sequences, and atom coordinates feature extraction, respectively. Detailed information of the above-mentioned strategies was summarized as follows.

### 2.3. Molecular graphic feature extraction

CNNs are highly successful graphic feature extractors, and are commonly developed with high accuracy by large training datasets. Given the utility of convolution kernels and DNNs, CNNs have played a critical role in image and video processing. Recently, molecular graph embedding has been used to model the relationships between chemical compounds (Sharma et al., 2021). To investigate the feasibility of molecular graphic presentation for odor category prediction, we used 4 types of effective CNNs, including the VGG-16, Resnet, Densnet, and Alexnet, as feature extractors for generating embedding from molecular images in the present study (Fig. 2a). Detailed structures for these CNNs have been previously presented (Krizhevsky et al., 2017). In the present study, we used pre-trained CNN models for odor category prediction to overcome the limitation of sample size.

### 2.4. Molecular sequence feature extraction

As an end-to-end supervised learning algorithm, graph neural networks (GNNs) have been widely applied for sequence embedding in various fields. Given that odorants can be described as molecular topology graphs using SMILES, we considered GNNs to be appropriate for molecular presentation. Broadly speaking, graph transformers are considered to be a powerful tool for handling molecular presentation through encoding via SMILES, which has been used to predict compound protein interactions, virtual screening, and molecular parameters (Warikoo et al., 2021).

Although the molecular graph transformer neural network (MGTNN) has strong potential for molecular modeling, deep learning models always require a large amount of labeled data for training. To overcome the above problems, we used a self-supervised graph transformer (GROVER) to obtain presentation information from the odorants for odor category prediction. A brief description of the GROVER is given in Fig. 2b. The pre-training architecture was mainly composed of two parts: i) a transformer-based neural network, and ii) a GNN for molecular structure extraction. The input of the model was an odorant graph presentation 𝒢 = (*V, E*), where *V* was the set of atoms and *E* was the set of bonds. Specifically, *v*_*i*_ ∈ *V* and *e*_*i,j*_ ∈ *E* were the *i*-th atom and bond between the *i*-th and *j*-th atom, respectively. The GNN was designed to embed extraction according to queries (**Q**), keys (**K**), and values (**V**) from the atoms in molecular graphs (𝒢*v*). The message transmission process of the GNN, as well as the neighborhood aggression between an atom (*v*) and its neighbors (𝒩 *v*) in an odorant (𝒢*v*), were adopted to iteratively (*L*) updatehidden states (**h***v*) for atom *v*, which can be written as:

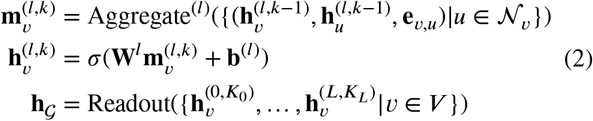

where 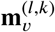 indicates the passing message for atom *v* under the *k*-th step of the *l*-th iteration. Here, we suppose each iteration (*l*) contains *K*_*l*_ steps. Aggregate^(*l*)^(⋅) is an aggregation function, which can be selected from the mean, max pooling, or graph attention mechanism. *σ*(⋅) is the activation function, and **h**_𝒢_ is the graph-level representation generated by a Readout operation. The resulting matrices (**Q, K, V**) were fed to the transformer module, which was composed of graph multi-head attention blocks:

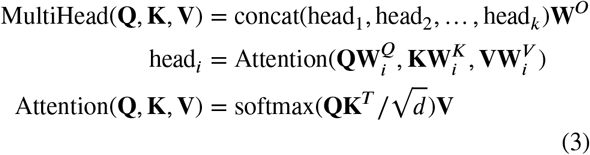

where 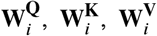 are the projection matrices of head_*i*_. *d* indicates the dimension of **q** and **k**.

The self-supervised learning tasks in the present study were assigned as contextual property prediction, and graph-level motifs, as well as molecular components, were used to predict links between both nodes and edges. In summary, we employed a pre-trained model, calibrated with 10 million molecules, as a molecular topology feature extractor in the present study. Instances of atom embedding (2048 dimensions) and bond embedding (2048 dimensions) generated by the above-mentioned procedure were used for odor category prediction. This simple strategy has been demonstrated to be a powerful method in terms of graph expression and structure information extraction. Details regarding graph transformers can be found elsewhere (Mao et al., 2021).

### 2.5. Atomic interaction embedding

Numerous studies have confirmed that atom interactions are crucial to odor perception (Fjaeldstad, 2018). Consequently, molecular features generated by atomic interactions may be feasible for odor category prediction. In the present study, we modeled interactions between atoms in an odorant molecule using a DNN-based model as a molecular feature extractor. A brief description of the process for the AINN is given in Fig. 2c. Formally, given an odorant 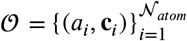, where *a*_*i*_ is the *i*-th atom type, **c**_*i*_ ∈ ℝ^2^*or*ℝ^3^ is the coordinate vector of the *i*-th atom, and *N*_*atom*_ is the total number of atoms for the odorant. To obtain an atom embedding description for the odorant, 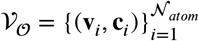, where **v**_*i*_ ∈ ℝ^*d*^ is the embedded vector for the *i*-th atom. The embedding dimensionality *d* is a hyper-parameter that must be assigned before training, and these atom embeddings were initialized randomly and optimized via backpropagation. To select the update strategy for the above-mentioned embeddings, we referred to the previous use of DNNs with common graph-structured datasets (Chen et al., 2020). Accordingly, the atom embeddings were updated as follows:

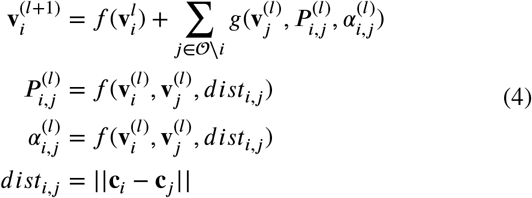

where *f*(⋅) and *g*(⋅) were the neural networks. 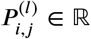 and 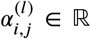 indicate the potential volume and interaction factor between the *i*-th and *j*-th atoms at the *l*-th hidden layer, respectively. *dist*_*i,j*_ ∈ ℝ was the Euclidean distance between the *i*-th and *j*-th atoms. Thus, the atom interaction embeddings for the odorant (**x**_𝒪_ ∈ ℝ^*d*^) could be calculated by:

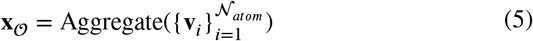

where Aggregate(⋅) was the aggregate function, which was mean pooling in the present study. As an option, we added a residual part to prevent the vanishing gradient problem in the DNN (res-AINN), which could be defined as follows:

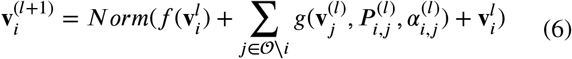

Finally, the odorant embeddings 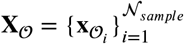 were selected as inputs for subsequent models. In this study, the hyper-parameters *L*, dimensions of embeddings *d*, learning rate *η*, and learning epochs were selected as 6, {32, 64, 128, 256, 512, 1024}, 0.001, and 200, respectively. We considered the feasibility of using molecular 2D and 3D coordinates, and discussed the embedding results. Detailed information regarding atom interaction embedding can be found in other publications (Tsubaki & Mizoguchi, 2019).

## 3. Results and discussion

### 3.1. Data analysis

We employed five different molecular structure representations, including odorant molecular parameters (MP), molecular fingerprints (FP), pre-trained molecular graphic embeddings, pre-trained molecular graph transformer embeddings, and atom interaction embeddings in the present study. First, we visualized the numeric vectors in lowdimensional space using Barnes-Hut t-distributed stochastic neighbor embedding (t-SNE) as an unsupervised statistical method. This method has been widely applied for high-dimensional data visualization. The t-SNE presentation of the odorants based on the above-mentioned vectors is illustrated in Fig. 3 and Fig. S1. As reported in previous studies, each odorant contained multi-odor category labels (Shang et al., 2017; Snitz et al., 2013; Kumar et al., 2015; Sánchez-Lengeling et al., 2019; Shang et al., 2021). Therefore, the distribution of odor categories was visualized using colors representing alpha values. Molecular graphic features extracted by Resnet (Fig. 3c) produced a better result than other molecular features because most odorants from the same odor category are clustered together. In contrast, odor cluster overlapping was observed more frequently in the t-SNE map generated from molecular fingerprints (Fig. 3a), molecular parameters (Fig. 3b), and molecular graph transformers (Fig. 3d). Molecular graphs generated from four combined types of pre-trained CNNs produced competitive results compared with other molecular descriptors (Fig. S1). This result demonstrates that odor categories are likely to be more strongly related to molecular graphs than other descriptors. Therefore, we inferred that an odor category identification model based on molecular graphic features would be superior.

**Figure 3:**
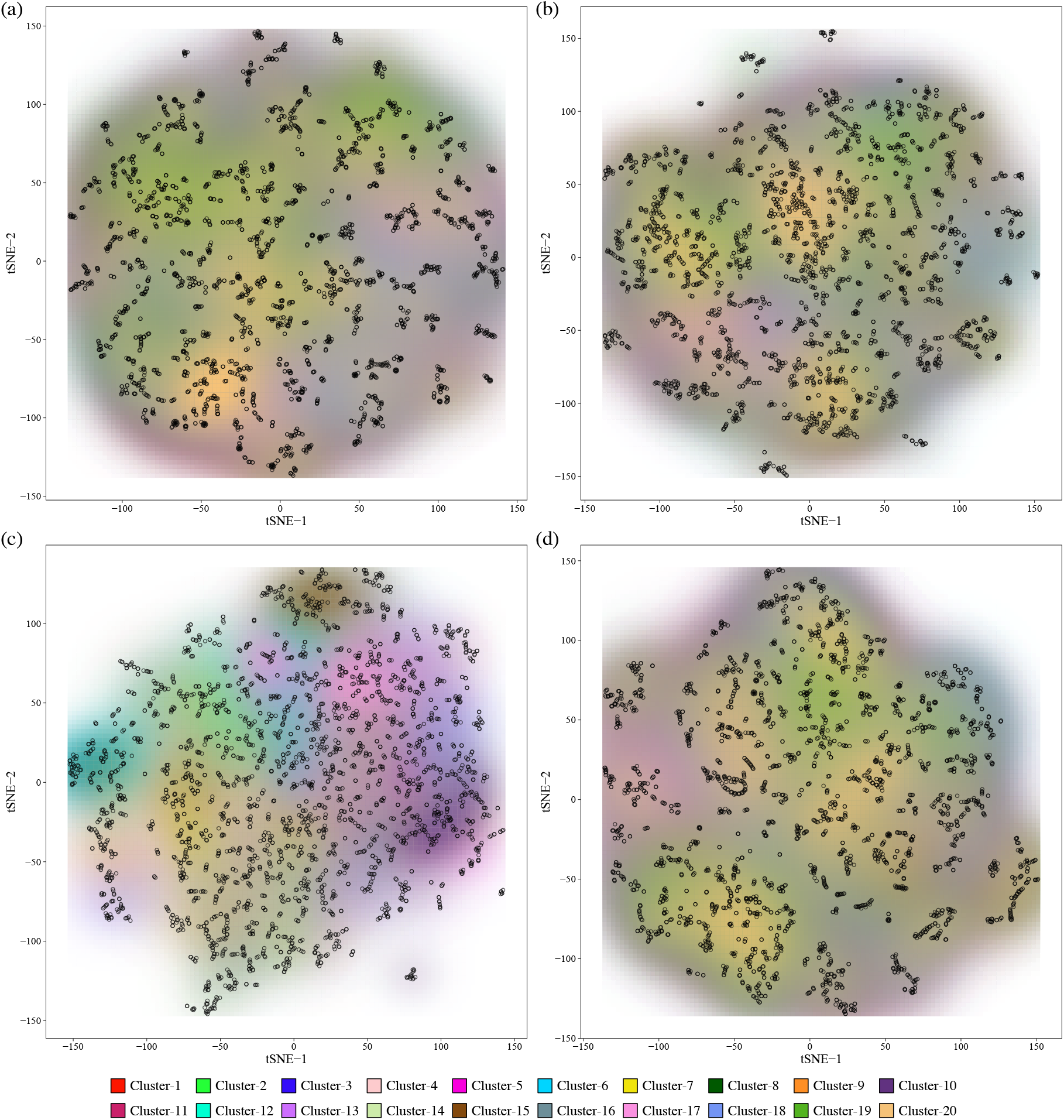
Odorant clustering using Barnes-Hut t-distributed stochastic neighbor embedding (t-SNE) based on (**a**) molecular fingerprints, (**b**) molecular parameters, and (**c**) molecular graphic features extracted via a pre-trained Resnet and (**d**) molecular graph transformer method based on the links between atoms and bonds. tSNE-1 and tSNE-2 were calculated using the t-SNE method. Each point indicates an odorant, colored according to its odor category labels, and the distributions of odor categories are given by the alpha values corresponding to the colors.

### 3.2. Molecular graphic CNN-based feature analysis

Fig. 4 and Table S2 summarize the performance metrics of the odor category identification model based on molecular graphic feature extraction. The details of model calibration, including the optimal epochs, training loss, and elapsed time, are illustrated in Fig. S2. The pre-trained ResNet with DNNs (6 hidden layers) performed significantly better than the other models, with the highest AUC (0.877±0.028, p<0.001) and F1-score (0.725±0.0278, p<0.001) on the test sets. It was followed by the DenseNet (4 hidden layers, AUC 0.876±0.029, F1-score 0.716±0.035), VGG (6 hidden layers, AUC 0.875±0.028, F1-score 0.716±0.033), and AlexNet (5 hidden layers, AUC 0.873±0.029, F1-score 0.723±0.033). The deep residual framework of the most successful model may have overcome the degradation problem that affects deep networks. In addition, the number of hidden layers in the DNN did not play a necessary role in tuning the pre-trained CNN model. To verify the abilities of the models for transfer learning, we compared the prediction performances of the CNNs depending on whether they were pretrained. These results are illustrated in Fig. 4. We found that the pre-trained models had significantly (p<0.001) higher accuracy compared with the models with un-trained CNNs. This indicates that CNNs could learn universal image feature extractors through training with a large dataset (ImageNet). This conclusion is supported by previous research (Liu et al., 2020).

**Figure 4:**
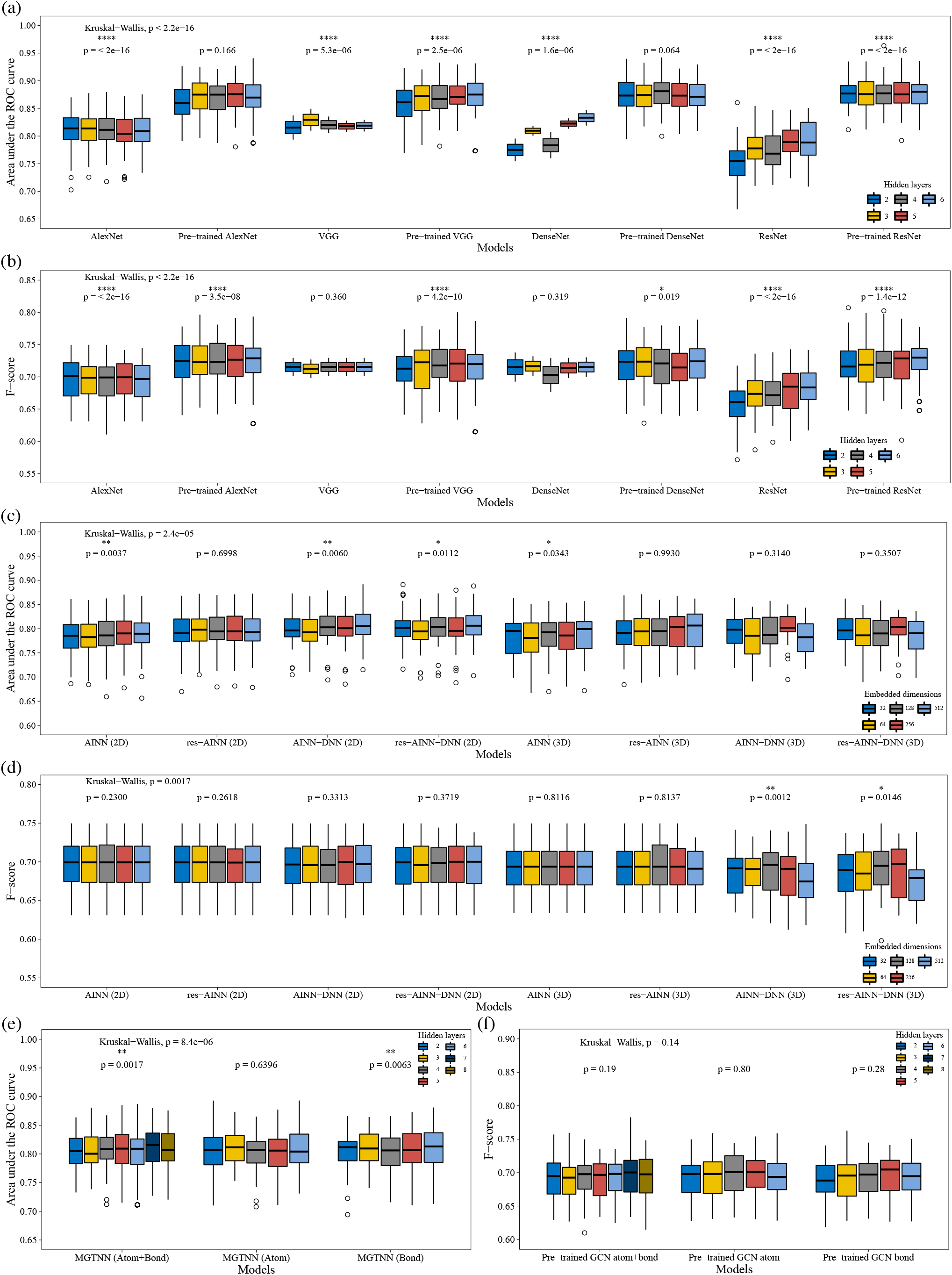
Identification performance of four molecular graphic convolution neural network (MG-CNN) models (**a, b**), molecular graph transformer neural network (MGTNN) models (**c, d**), and atom interaction neural network (AINN) models (**e, f**). Results were evaluated using the nonparametric Wilcoxon signed-rank test.

### 3.3. Molecular graph transformer-based feature analysis

A summary of the identification accuracy of the MGTNN models is given in Fig. 4 and Table S3. The optimal training epoch, loss, and elapsed time for the MGTNN models are presented in Fig. S3. When the selected atom and bond embeddings were included with 7 hidden layers, the MGTNN model had the highest AUC (0.813±0.035) and F1-score (0.696±0.032) in the test set. In addition, the AUC values for the models independently trained via atom or bond embeddings were 0.812±0.031 and 0.810±0.030, respectively. However, the data were not sufficient to conclude that considering atoms and bonds together produced a significantly more accurate result than when they were included individually (p>0.001).

### 3.4. Atom interaction-based feature analysis

Fig. 4 and Table S4 compare odor sensory category identification according to the molecular features extracted by AINNs. The results indicated that the AINN-DNN model (2D, 512 embedded dimensions) had the highest identification performance in terms of the AUC and F1-score, which were 0.807±0.035 (p<0.01) and 0.696±0.023, respectively. However, we cannot claim that the molecular 2D coordinates were better for identification than the 3D coordinates because the analyses for both had a high p-value. Furthermore, the models with residual modules did not exhibit a significant increase, likely because the vanishing gradient is not the critical obstacle limiting AINN performance. In addition, the dimension of atom embedding vectors did not have a significant effect on the accuracy of odor category identification. The optimal training epoch, loss, and elapsed time for AINN models are listed in Fig. S4. We found that the modeling time for 2D coordinates was significantly smaller than that for 3D coordinates (p<0.001). This suggests that the presented AINN models do not need spatial embedding for odor sensory identification.

### 3.5. Performance comparison

To identify the model with the best comprehensive performance for odor category identification, we compared the five types of models in terms of performance metrics, as presented in Fig. 5 and Table 1. The predicted accuracies for each odor sensory category are summarized in Fig. S6-S10. The results confirmed that the model trained using molecular graphic features extracted via a pre-trained ResNet had significantly better performance than the other models (AUC 0.877±0.028, F1-score 0.726±0.028, p<0.0001), followed by the AINN-DNN (AUC 0.807±0.035, F1-score 0.696±0.030), MPs (AUC 0.806±0.033, F1-score 0.689±0.031), MGTNN (AUC 0.804±0.028, F1-score 0.692±0.029), and FPs (AUC 0.796±0.036, F1-score 0.688±0.033). This ranking could likely be explained by the high correlation between the olfactory sensory information and the molecular graphic features of the odorants compared with the other molecular descriptors. We found that the AINN-DNN model had the highest precision (0.861±0.038, p<0.0001). Fig. S5 illustrates the optimal training epoch, loss, and elapsed time for the above-mentioned models. We confirmed that although more epochs were needed to train the ResNet models, the training time was shorter than that for the AINN-DNN and MGTNN models. The fast convergence speed could contribute to the transfer-learning mechanism. Although the number of parameters was abundant for the ResNet model, we did not train these parameters, but instead used those from the pre-trained models. The pre-trained models could overcome the limitation of insufficient samples for training DNN models. A similar conclusion was found previously (Pezzotti et al., 2017). In summary, we suggest that an end-to-end DNN with molecular graphic features extracted via a pre-trained ResNet is an optimal model for predicting the sensory categories of odorants.

**Figure 5:**
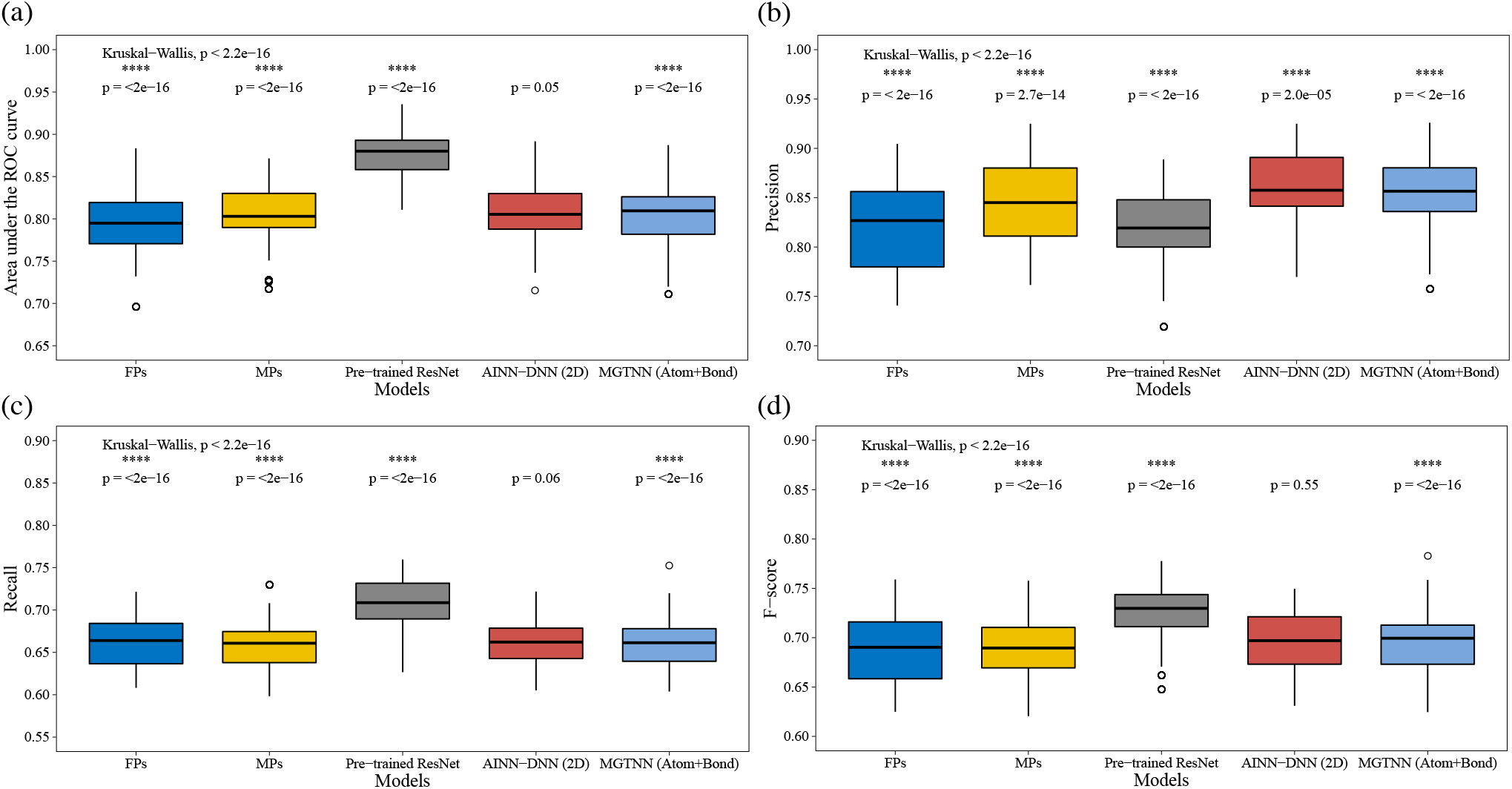
Identification accuracies of DNN models using molecular features extracted via FP-, MP-, MG-CNN-, MGTNN-, and AINN-based DNN models. The data were evaluated using the nonparametric Wilcoxon signed-rank test.

**Table 1.** Odor sensory category identification accuracy comparison of DNN models using multi-type of odorant structure features.

| Model name | AUC-ROC | Precision | Recall | F1-score |
| --- | --- | --- | --- | --- |
| FP-based | 0.796±0.036 | 0.82±0.046 | 0.662±0.029 | 0.688±0.033 |
| MP-based | 0.806±0.033 | 0.843±0.039 | 0.658±0.028 | 0.689±0.031 |
| MG-CNN-based | 0.877±0.028 | 0.822±0.037 | 0.71±0.028 | 0.726±0.028 |
| MGT-NN-based | 0.804±0.036 | 0.855±0.036 | 0.659±0.025 | 0.692±0.029 |
| AINN-based | 0.807±0.035 | 0.861±0.038 | 0.662±0.027 | 0.696±0.03 |

## 3.6. Discussion

The accurate and effective prediction of odor sensory categories is vital for developing machine-learning-based GCO. To develop an olfaction-based sensory system, we need not only bio-sensors to encode odorants (odor receptor imitation), but also a brain-like odor signal decoding algorithm. Although many studies have examined SORs, most have focused on predicting ODs (Snitz et al., 2013; Debnath & Nakamoto, 2020; Loetsch et al., 2019; Sharma et al., 2021; Kumar et al., 2015). Even though most ODs can be predicted, infrequent ODs were difficult to be identified. For example, Snitz proposed a mostly perfect result in predicting 64 smell percepts with 100 % precision and 102 smells with 90.35 %, but infrequent smells, such as almond, apricot, and chocolate, had been found to have poor prediction performance (Snitz et al., 2013). This could be explained by the extreme imbalance in the data distribution, as well as the insufficient number of training samples. Unlike the above-mentioned studies, we want to find useful odorant structure features for odor sensory categories identification.

Here, we focused on establishing relationships between molecular features and odor sensory categories via an end-to-end learning strategy, which is expected to play a decoding role in bio-olfaction. The MPs and FPs in the present study had poor performance, indicating that focusing solely on physiology-chemical parameters could result in the loss of some critical information related to olfaction. In contrast to relying on tabular features, molecular graph CNNs-based features would be more appropriate for learning useful odor sensory expression. We also considered a transfer learning strategy for dealing with the problem of insufficient training samples. Our results confirm that pre-trained CNNs, combined with a ‘vanilla’ DNN, can effectively establish relationships between molecular features and odor sensory categories. Furthermore, our data suggest that molecular graphic features are optimal for describing odorant protein interactions according to human olfaction. Existing GCO methods have focused on just 8 ODs in one olfaction sensory evaluation task, as limited by the odor memory of the assessors (Brattoli et al., 2013; Vene et al., 2013). Odor analysis precision is also limited by their odor memory. Therefore, the proposed model can apply reliable references for human panelists to reduce training costs.

This study has several limitations. First, more attention should be focused on atom interaction-based embeddings, although the AINN in the present study had poor performance. Biological studies have indicated that atom interactions play a critical role in mammal olfaction (Mantel et al., 2022). The poor accuracy of the AINN was likely caused by insufficient odorants and an inappropriate modeling approach. Moreover, we did not consider the electronic interactions between atoms, which may be suitable for olfaction sensory encoding. Self-supervised strategies combined with proper modeling techniques and trained with abundant molecules merit further investigation. Furthermore, synergism, odor neutralization, and the predictability of a fragrance mixture have still not been quantified (Hudon et al., 2000). In the present study, we focus on single odor molecule smell perception prediction, which is not appropriate for modeling odor synergism and neutralization. For fragrance mixture prediction, mass spectral would be feasible for model calibration. In the future, we plan to attempt to improve our framework for molecular structure feature extraction using other algorithms, and to try to explore the feasibility of metric modeling using Riemannian manifolds, such as the Grassmann or symmetric positive definite manifold (Tang et al., 2021). We expect that it will be difficult to find a reasonable algorithm when performing metric learning in Riemannian space. However, this is an interesting problem for future investigation. In addition, data fusion would also be an effective strategy for increasing the accuracy of odor category identification models.

## 4. Conclusions

In summary, we utilized a DNN-based multi-label classifier for odor sensory category identification using various molecular features. Specifically, we examined the possibility of predicting odor categories based on molecular parameters, fingerprints, and graphics, as well as graph attention network embedding and atom interactions. Our results indicated that molecular 2D graphic data were strongly related to sensory information about olfaction. Extensive experiments confirmed that a ‘vanilla’ DNN with molecular graphic features, extracted via ResNet, was optimal for odor perception category identification. We anticipate that transfer learning is a viable and powerful technique for modeling the relationships between molecular structures and odor perception categories. Our proposed approach could be applied in the development of AI-based odor sensors. We believe that this study is among the first to examine the importance of molecular graphic features when establishing relational models between molecular structures and odor sensory categories. Our approach may not only serve as a realistic solution for introducing AI into olfactometry, but may also offer a novel auxiliary tool in the existing GCO by suggesting possible odor categories to the human assessors.

## Acknowledgements

This research was supported by China Postdoctoral Science Foundation (2021M703399), the National Nature Science Foundation of China (No. 61803369 and 61801400), and JSPS KAKENHI Grant (No. 18H03782).

## Appendix A.

Supplementary data

Supplementary data associated with this article can be found, in the online version, at http://dx.doi.org/.

## CRediT authorship contribution statement

**Liang Shang:** Conceptualization, Methodology, Experiment, Writing. **Chuanjun Liu:** Conceptualization, Method, Supervision, Reviewing. **Fengzhen Tang:** Supervision, Reviewing. **Bin Chen:** Supervision, Reviewing. **Lianqing Liu:** Reviewing. **Kenshi Hayashi:** Reviewing.

