## Supplementary material for "Artificial intelligence-based gas chromatography-olfactometry for sensory evaluation of key compounds in food ingredients": SI

### Contents

**Supplementary Figures and Tables** **3**

Supplementary Figures . . . . . . . . . . . . . . . . . . . . . . . . . . . . . . 3

Figure S1. Odorant clustering by using Barnes-Hut t-distributed stochastic neighbor embedding. . . . . . . . . . . . . . . . . . . . . . . . . . 3

Figure S2. Optimal training epoch, loss and elapsed time for MG-CNN models. 5

Figure S3. Optimal training epoch, loss and elapsed time for MGTNN models. 6

Figure S4. Optimal training epoch, loss and elapsed time for AINN models. . 7

### Supplementary Figures and Tables

#### Supplementary Figures

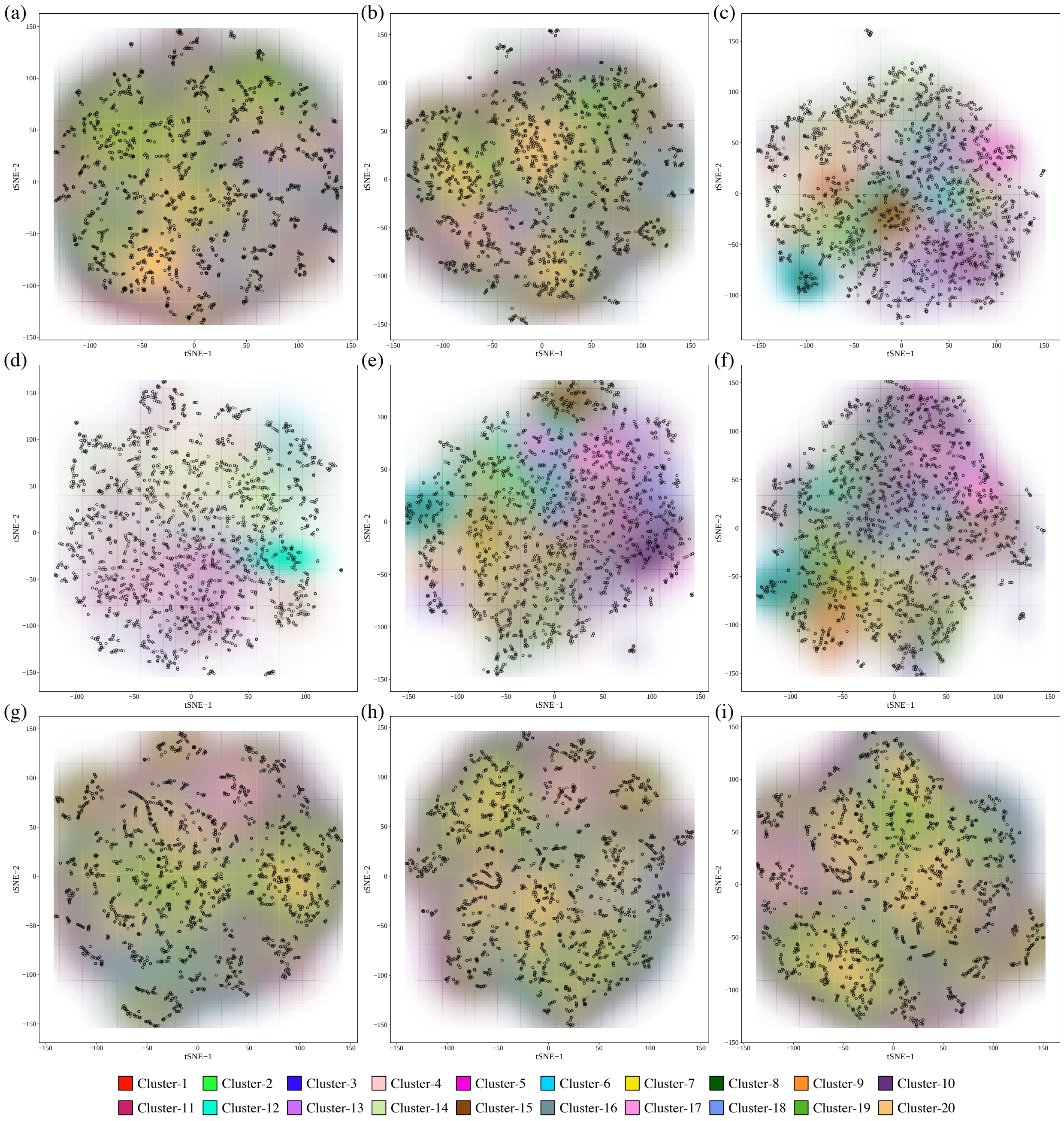

Figure S1: Odorant clustering by using Barnes-Hut t-distributed stochastic neighbor embedding (t-SNE) based on (**a**) molecular finger prints, (**b**) molecular parameters, molecular graphic features extracted by pre-trained (**c**) Alexnet, (**d**) Densenet, (**e**) Restnet and (**f**) VGGs, molecular graph transformer extraction based atom (**g**), bond (**h**) and atom+bond (**i**). tSNE-1 and tSNE-2 are the presentations calculated by t-SNE method. Each point indicates an odorant which colored by its odor category labels, and the distribution of odor categories are presented by the alpha of colors.

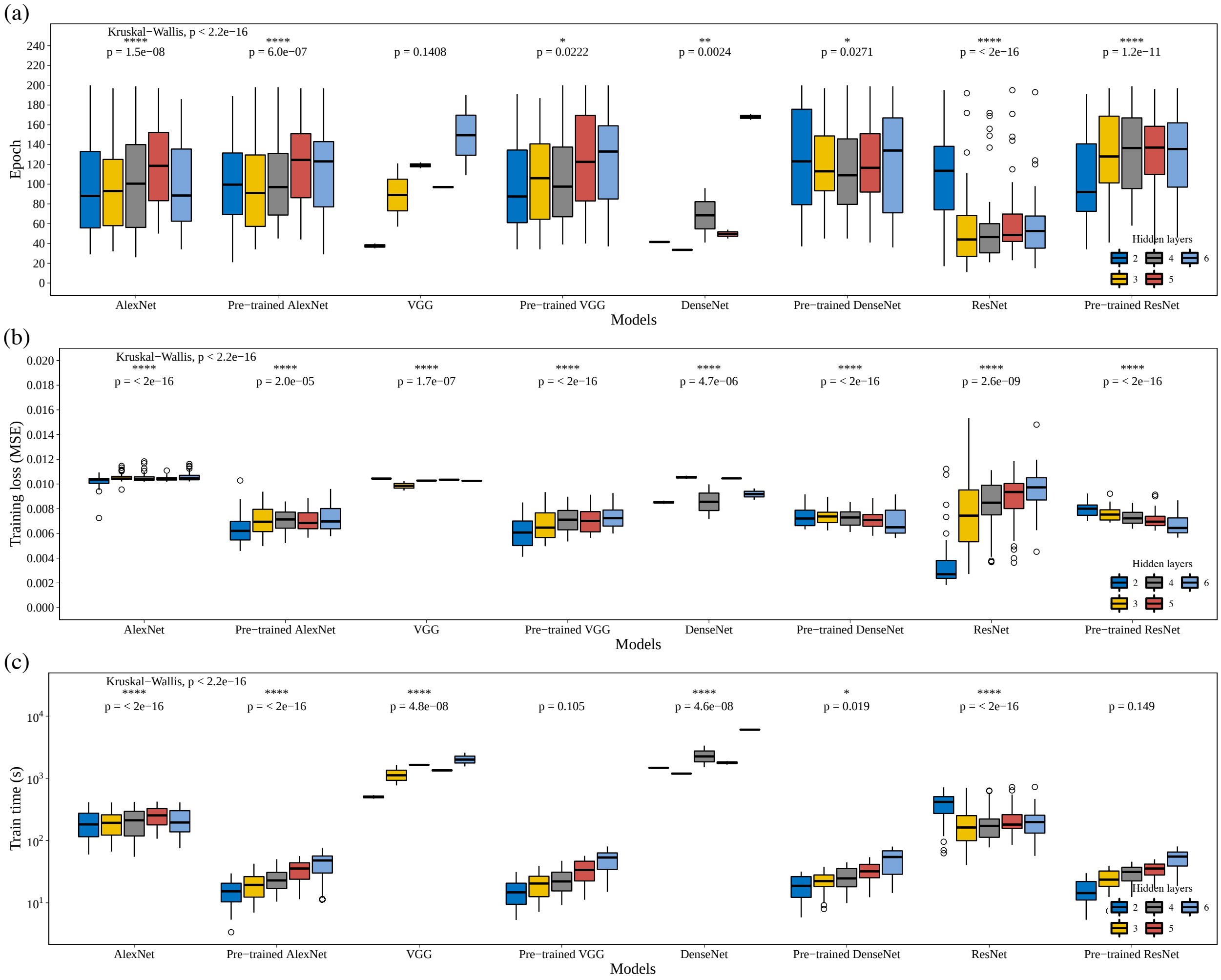

Figure S2: Optimal training epoch (**a**), loss (**b**) and elapsed time (**c**) for MG-CNN models.

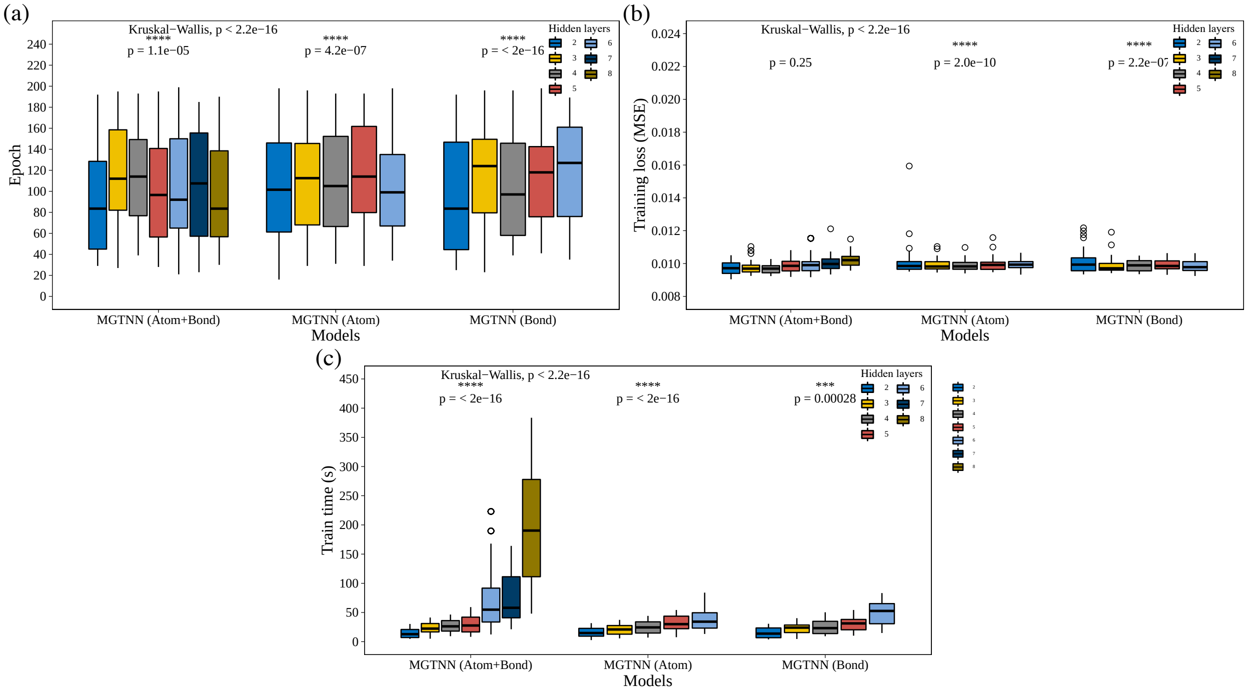

Figure S3: Optimal training epoch (**a**), loss (**b**) and elapsed time (**c**) for MGTNN models.

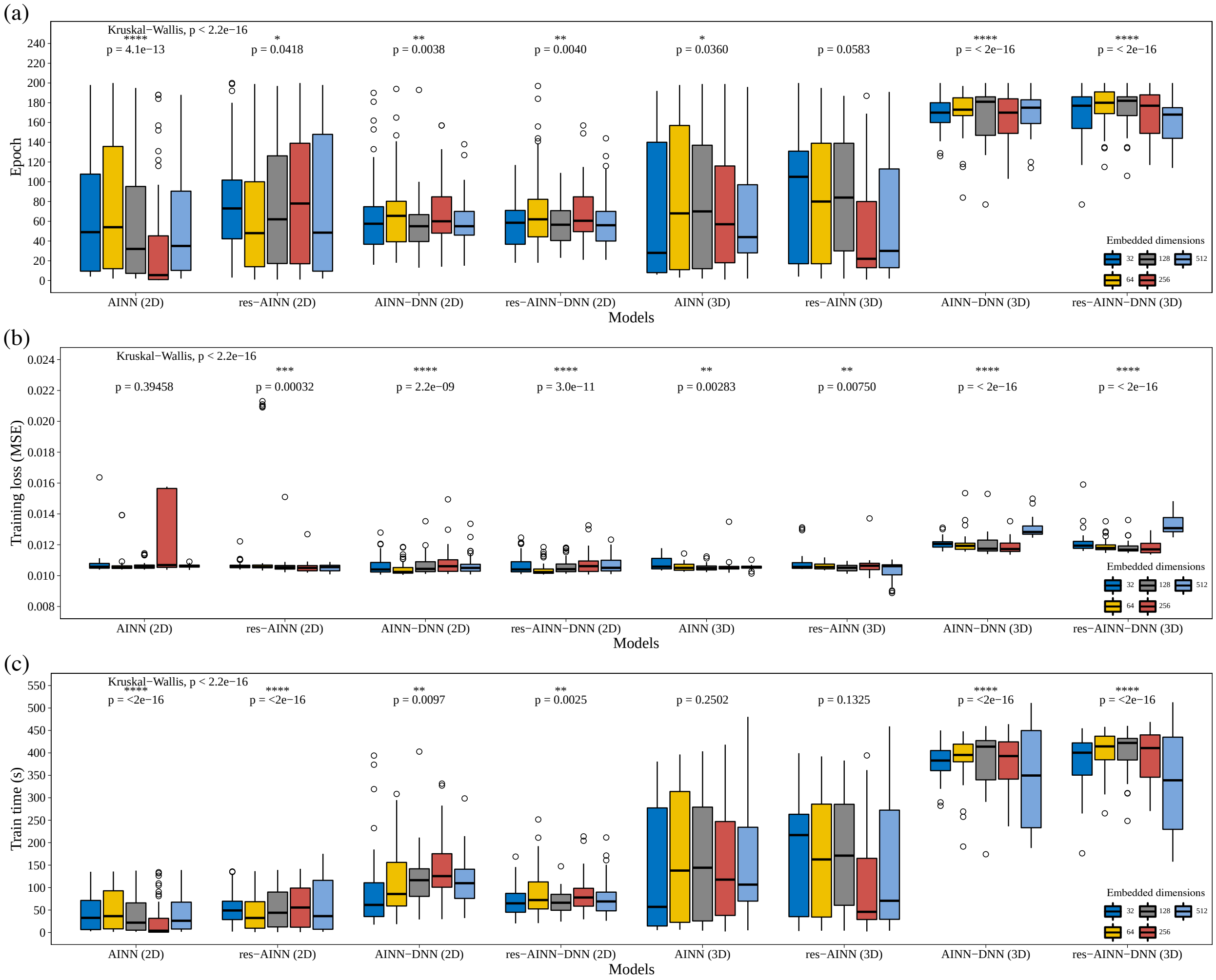

Figure S4: Optimal training epoch (**a**), loss (**b**) and elapsed time (**c**) for AINN models.

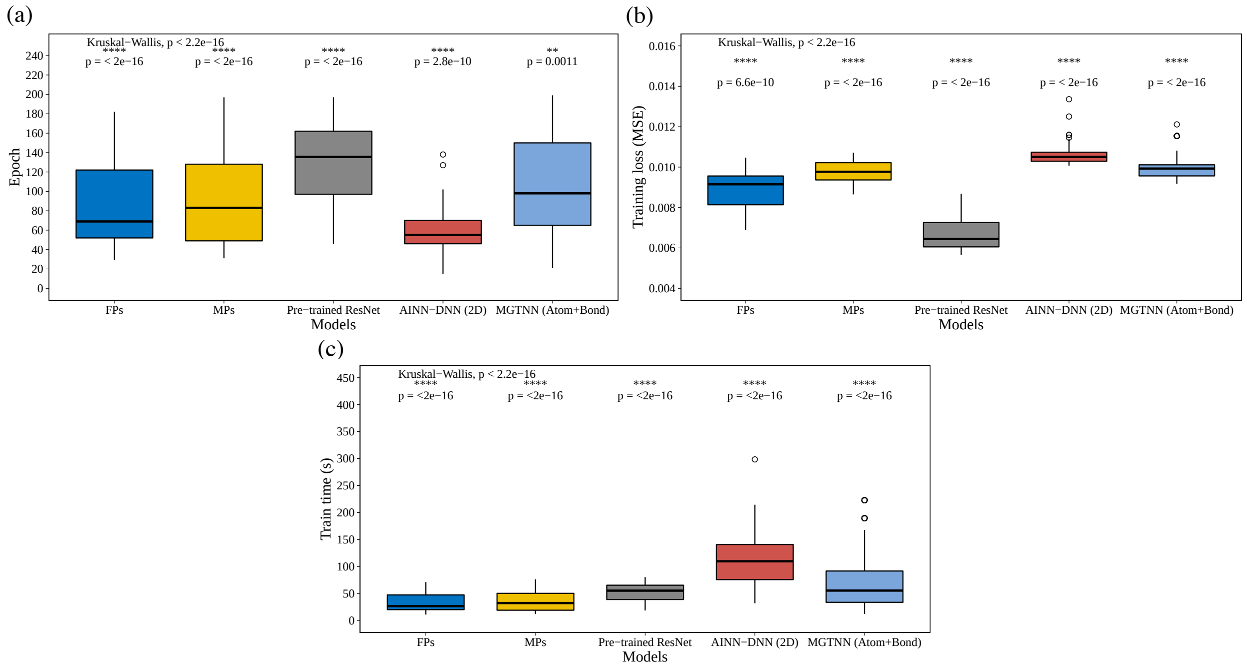

Figure S5: Training epoch (**a**), loss (**b**) and elapsed time (**c**) for optimal models.

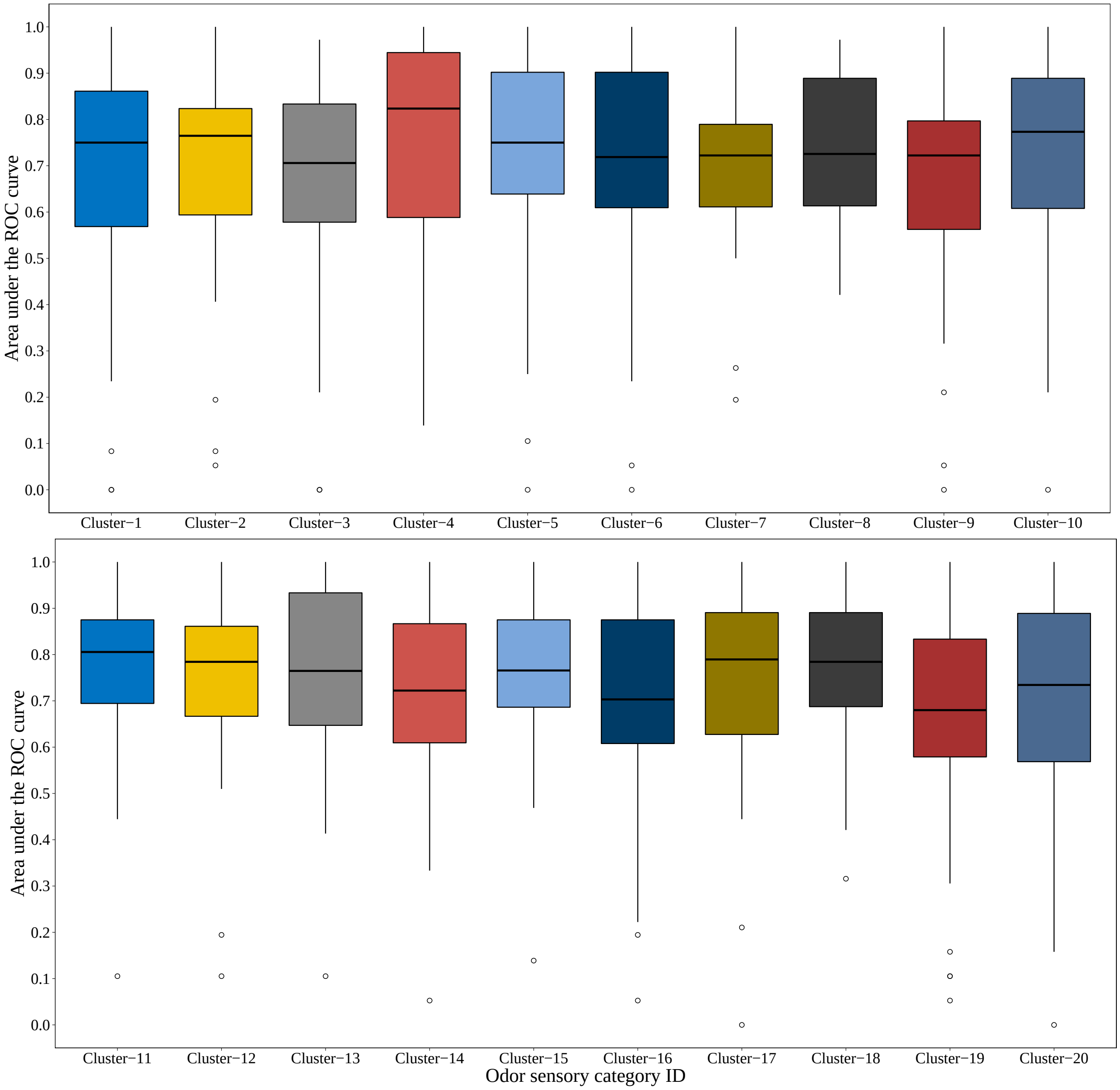

(a) Area under the ROC curve for molecular finger print based models.

Figure S6: Comparison of odor clusters identification for area under the ROC (**a**), precision (**b**), recall (**c**) and F-score (**d**) by molecular finger print based models.

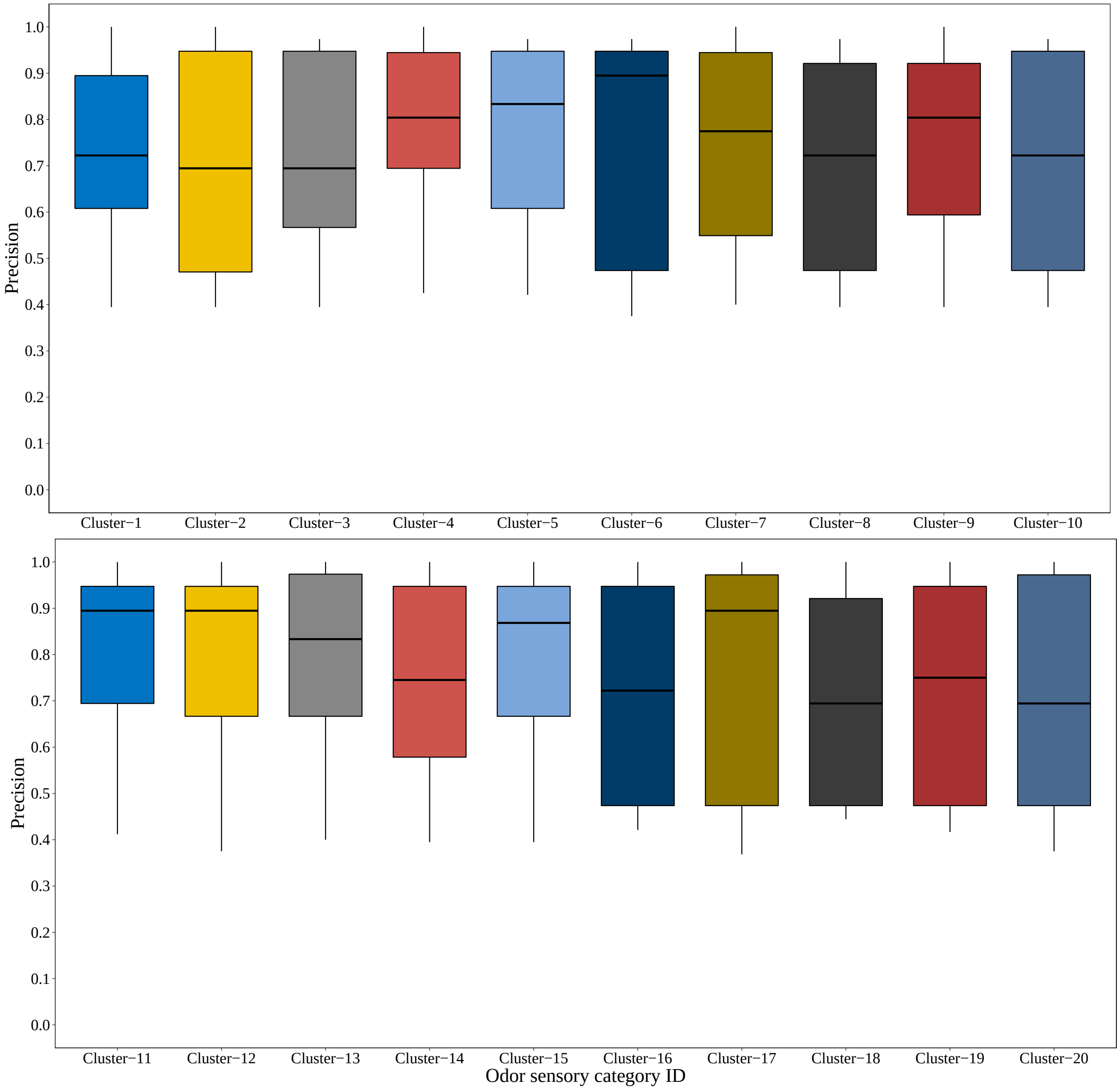

(b) Precision for molecular finger print based models.

Figure S6: Comparison of odor clusters identification for area under the ROC (**a**), precision (**b**), recall (**c**) and F-score (**d**) by molecular finger print based models.

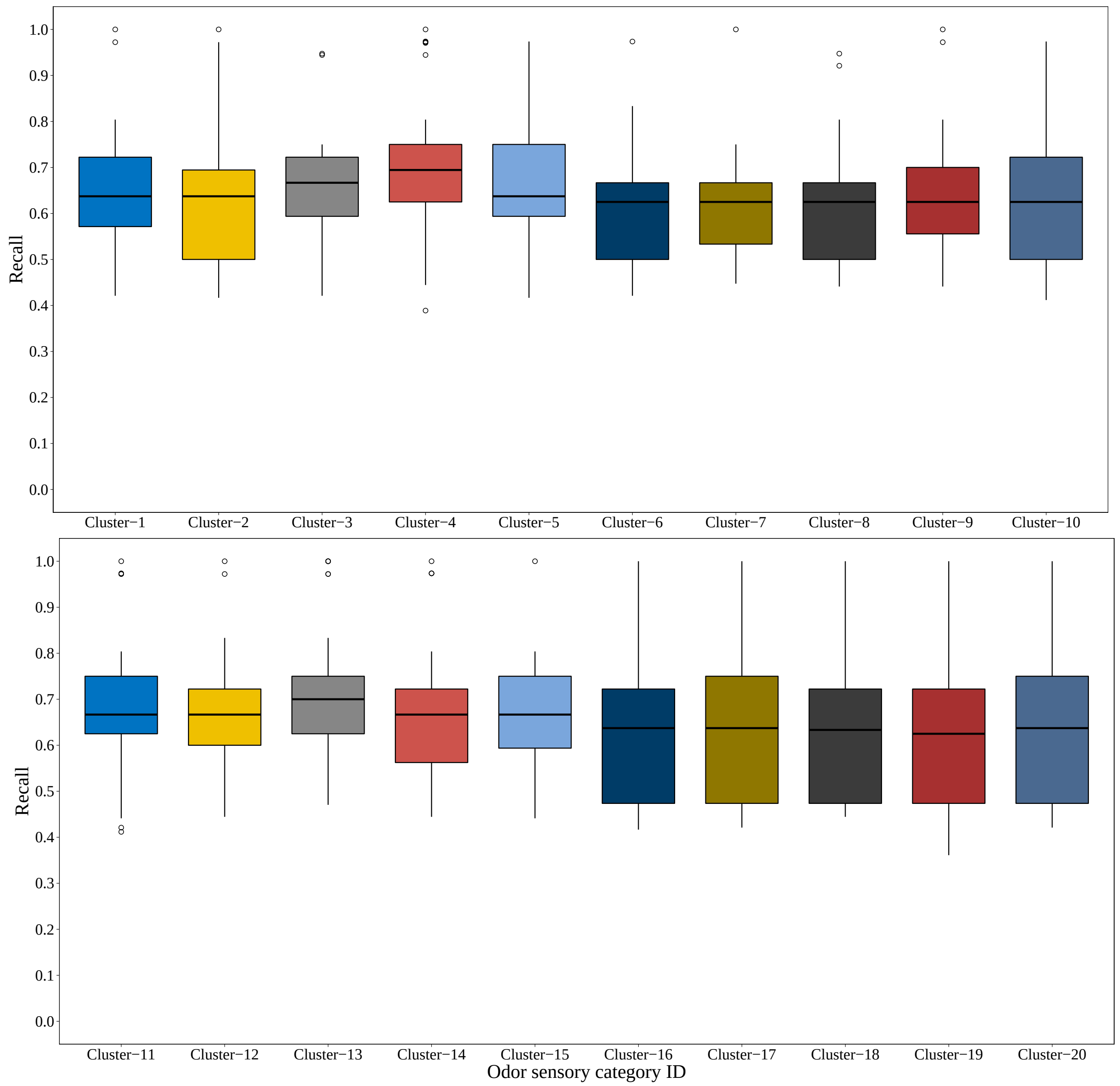

(c) Recall for molecular finger print based models.

Figure S6: Comparison of odor clusters identification for area under the ROC (**a**), precision (**b**), recall (**c**) and F-score (**d**) by molecular finger print based models.

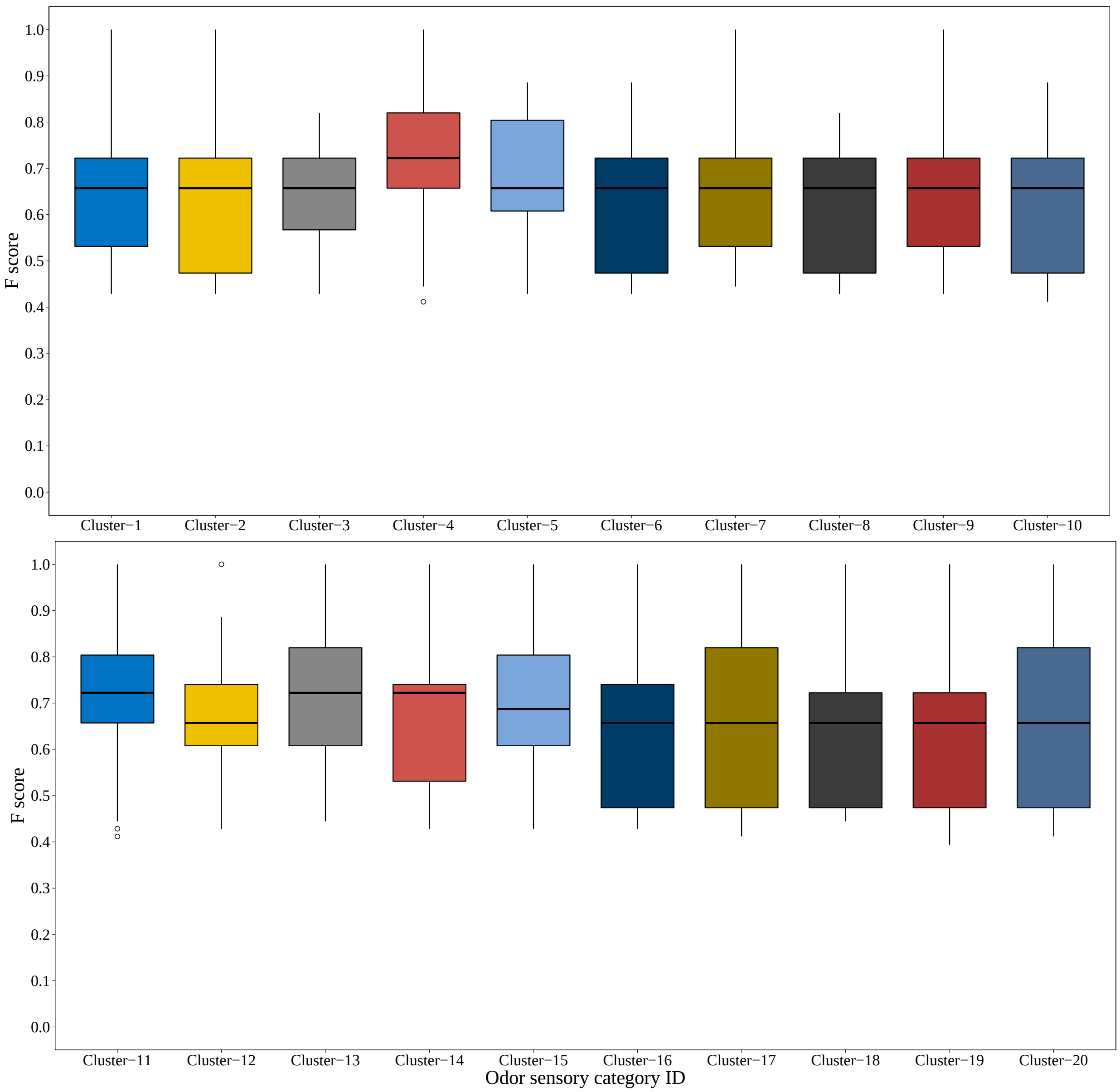

(d) F-score for molecular finger print based models.

Figure S6: Comparison of odor clusters identification for area under the ROC (**a**), precision (**b**), recall (**c**) and F-score (**d**) by molecular finger print based models.

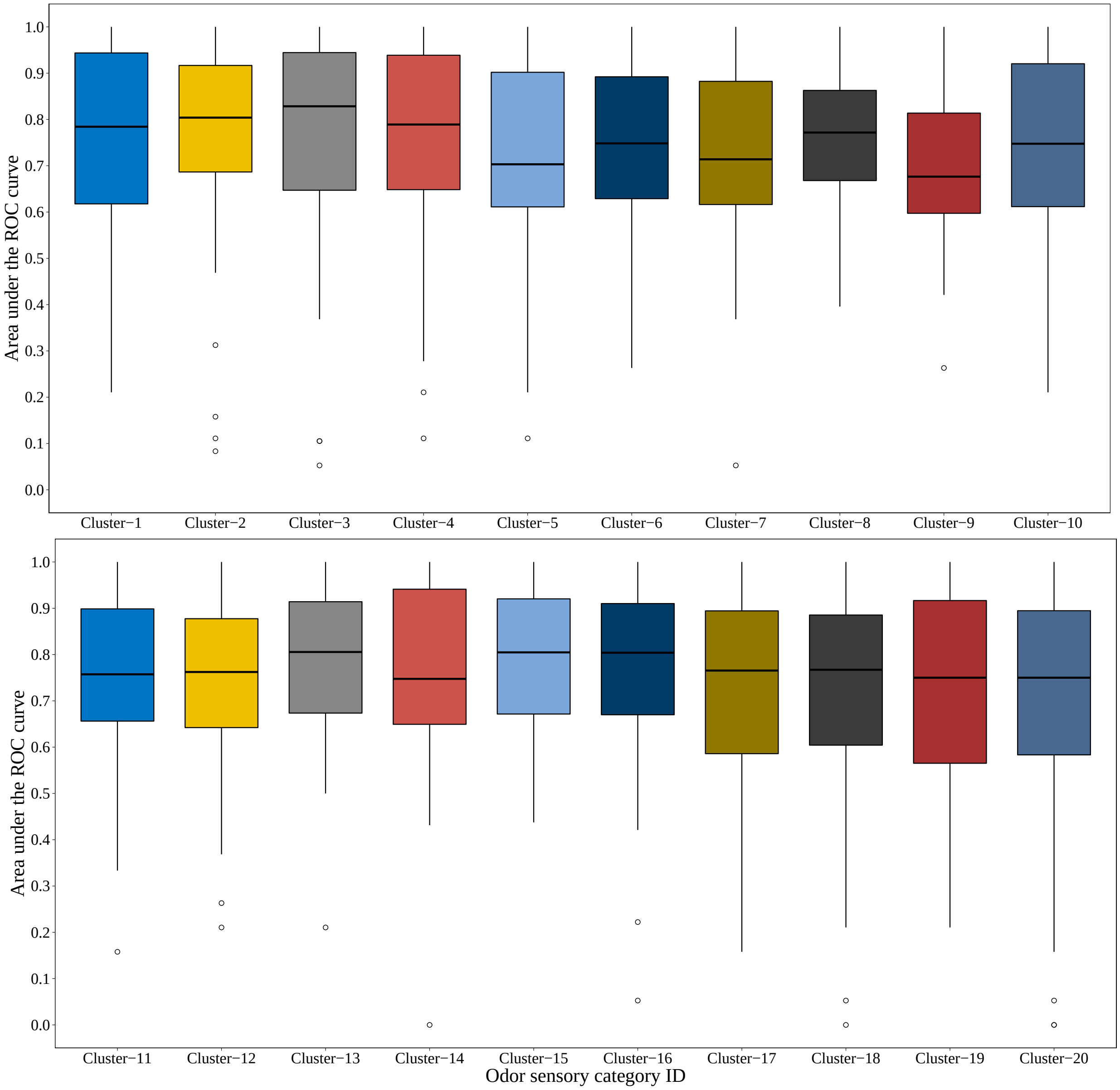

(a) Area under the ROC curve for molecular molecular parameter based models.

Figure S7: Comparison of odor clusters identification for area under the ROC (**a**), precision (**b**), recall (**c**) and F-score (**d**) by molecular parameter based models.

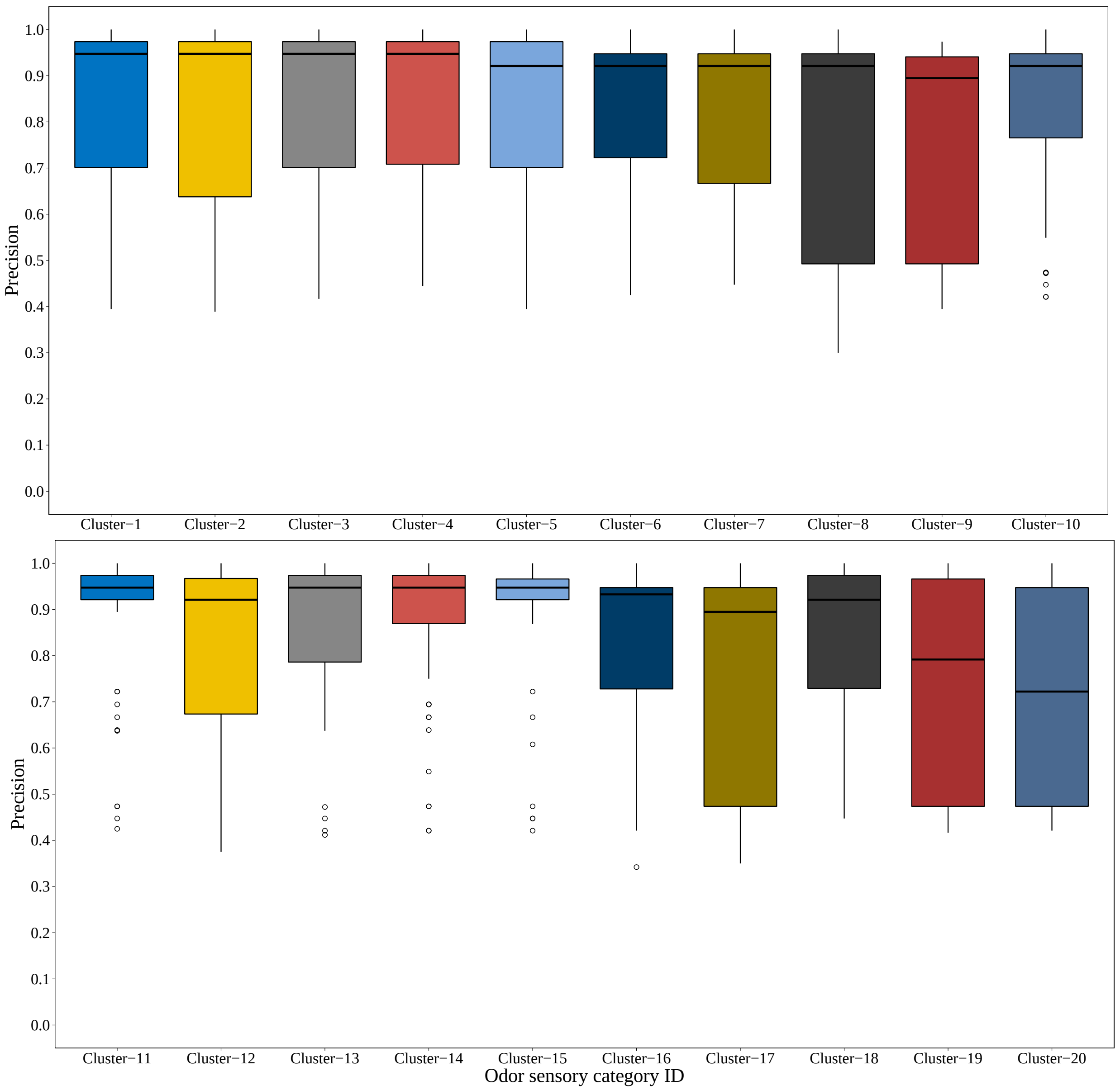

(b) Precision for molecular parameter based models.

Figure S7: Comparison of odor clusters identification for area under the ROC (**a**), precision (**b**), recall (**c**) and F-score (**d**) by molecular parameter based models.

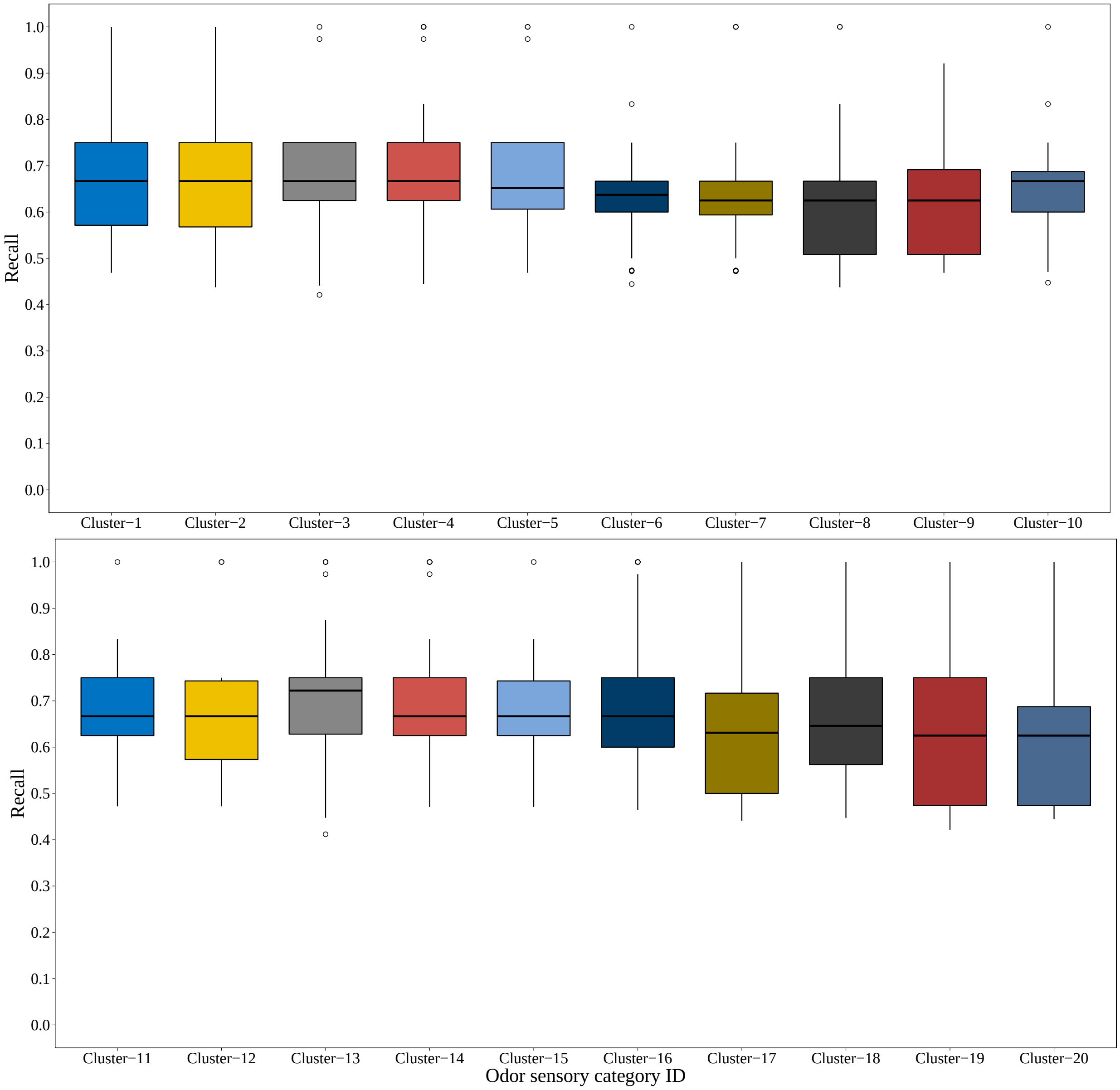

(c) Recall for molecular parameter based models.

Figure S7: Comparison of odor clusters identification for area under the ROC (**a**), precision (**b**), recall (**c**) and F-score (**d**) by molecular parameter based models.

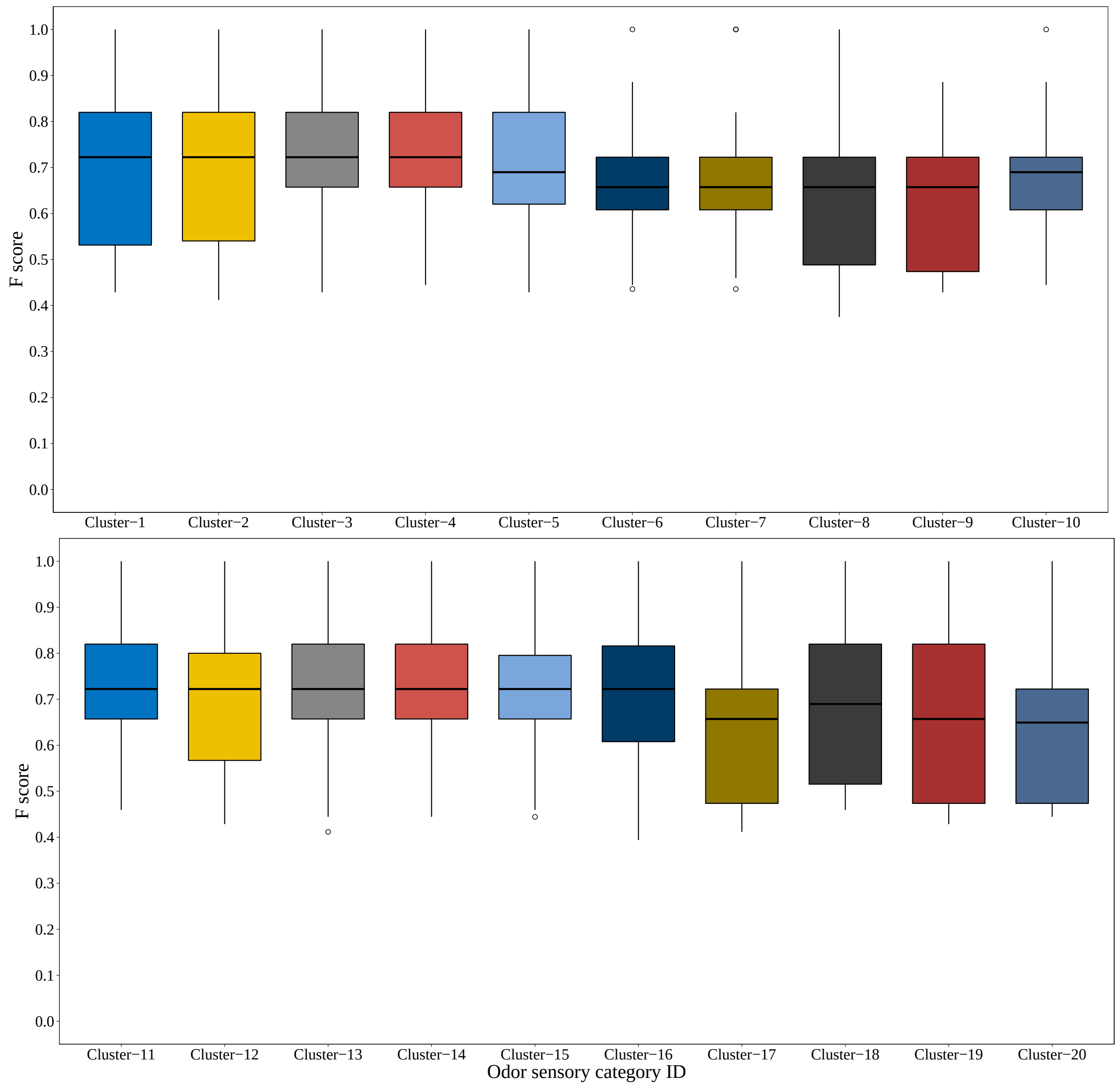

(d) F-score for molecular parameter based models.

Figure S7: Comparison of odor clusters identification for area under the ROC (**a**), precision (**b**), recall (**c**) and F-score (**d**) by molecular parameter based models.

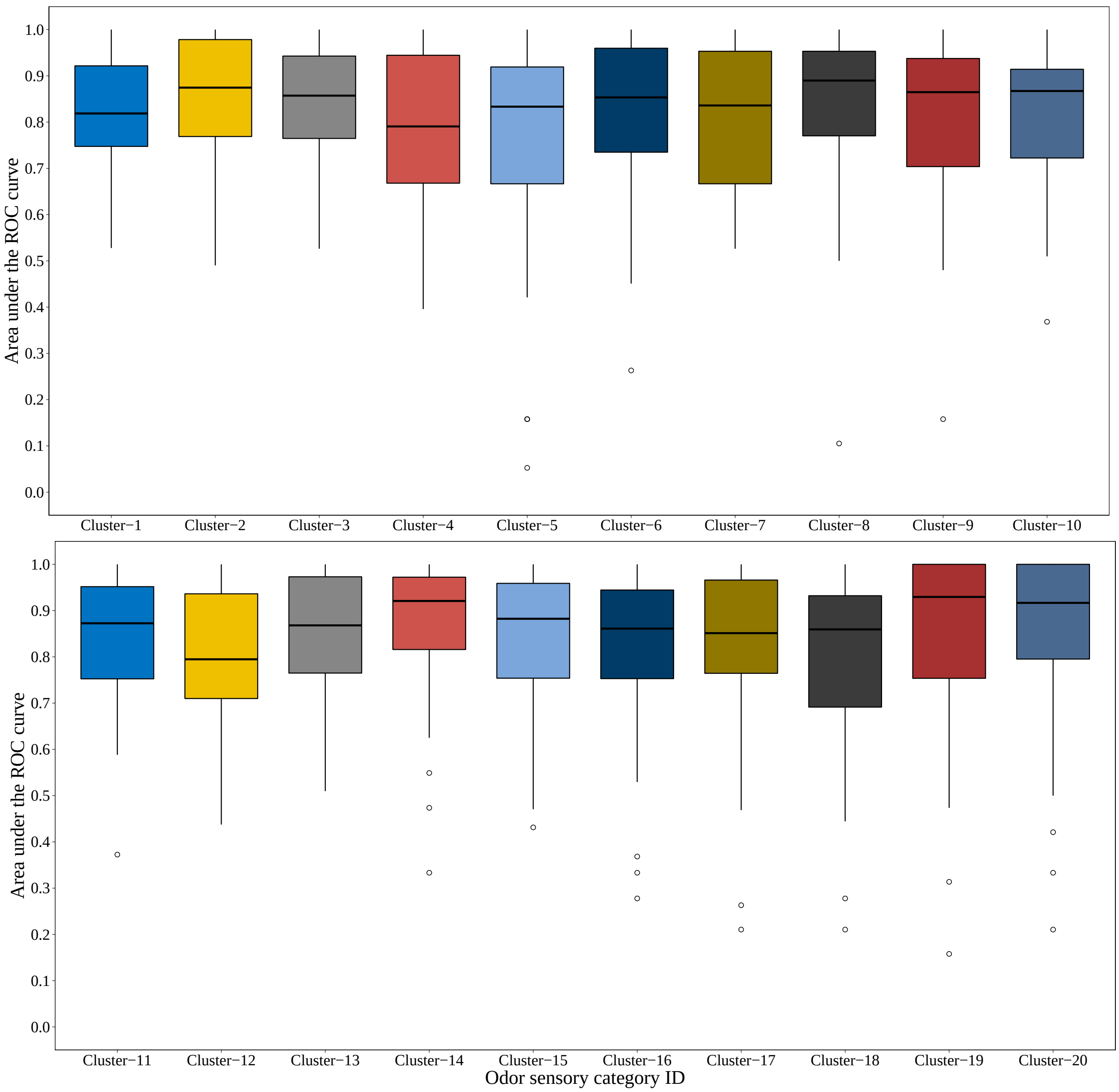

(a) Area under the ROC curve for MG-CNN models.

Figure S8: Comparison of odor clusters identification for area under the ROC (**a**), precision (**b**), recall (**c**) and F-score (**d**) by MG-CNN models.

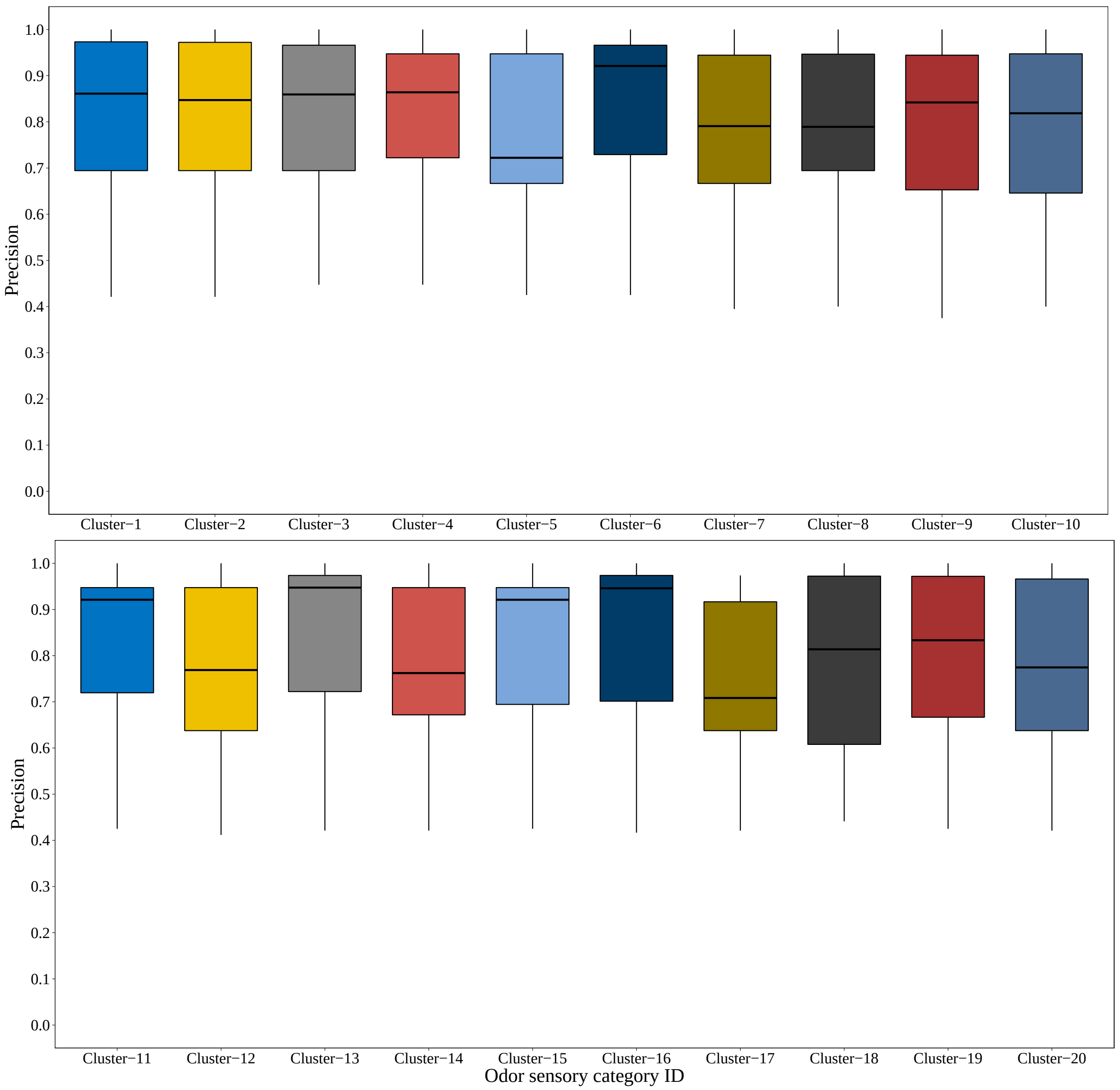

(b) Precision for MG-CNN models.

Figure S8: Comparison of odor clusters identification for area under the ROC (**a**), precision (**b**), recall (**c**) and F-score (**d**) by MG-CNN models.

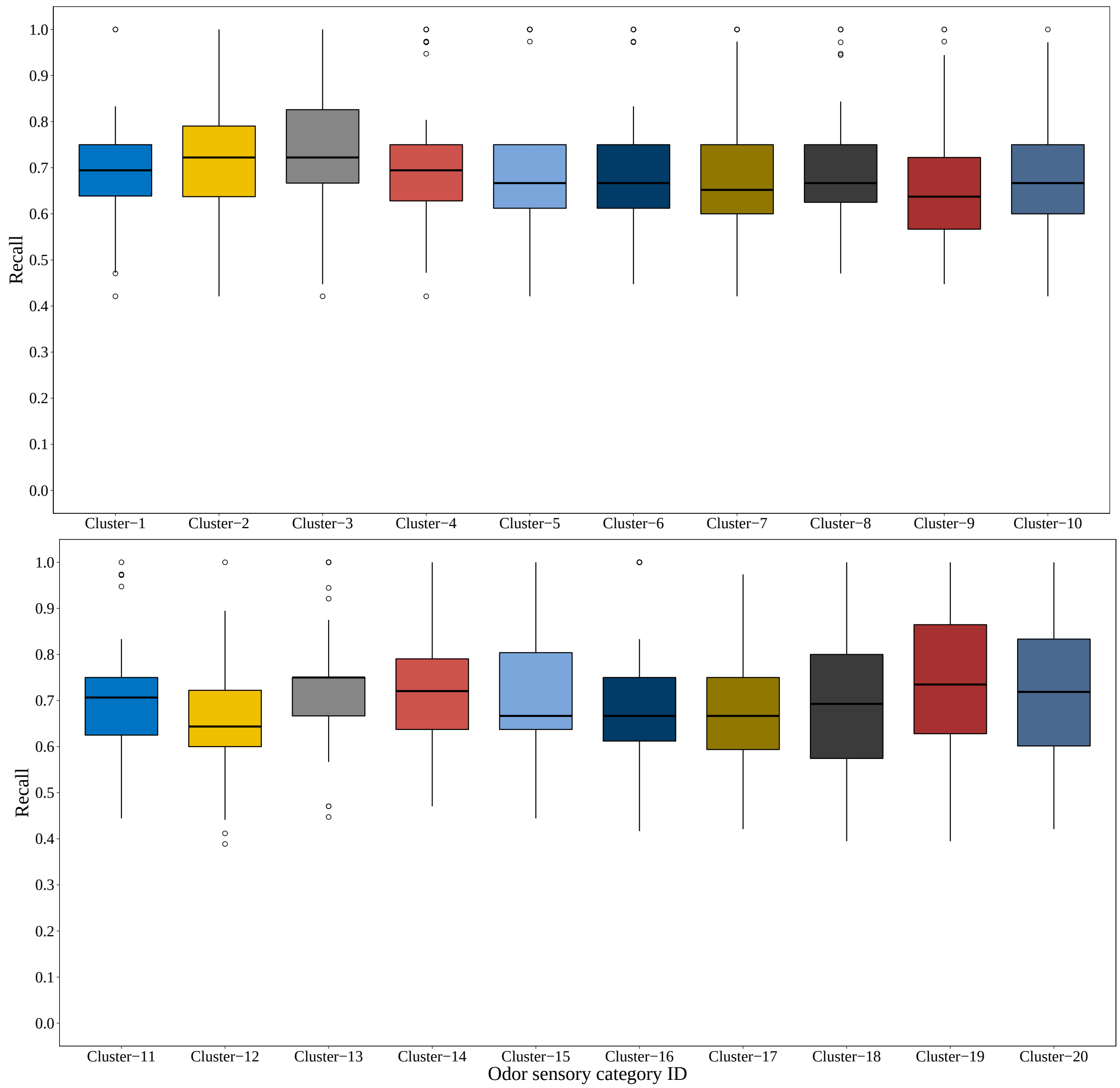

(c) Recall for MG-CNN models.

Figure S8: Comparison of odor clusters identification for area under the ROC (**a**), precision (**b**), recall (**c**) and F-score (**d**) by MG-CNN models.

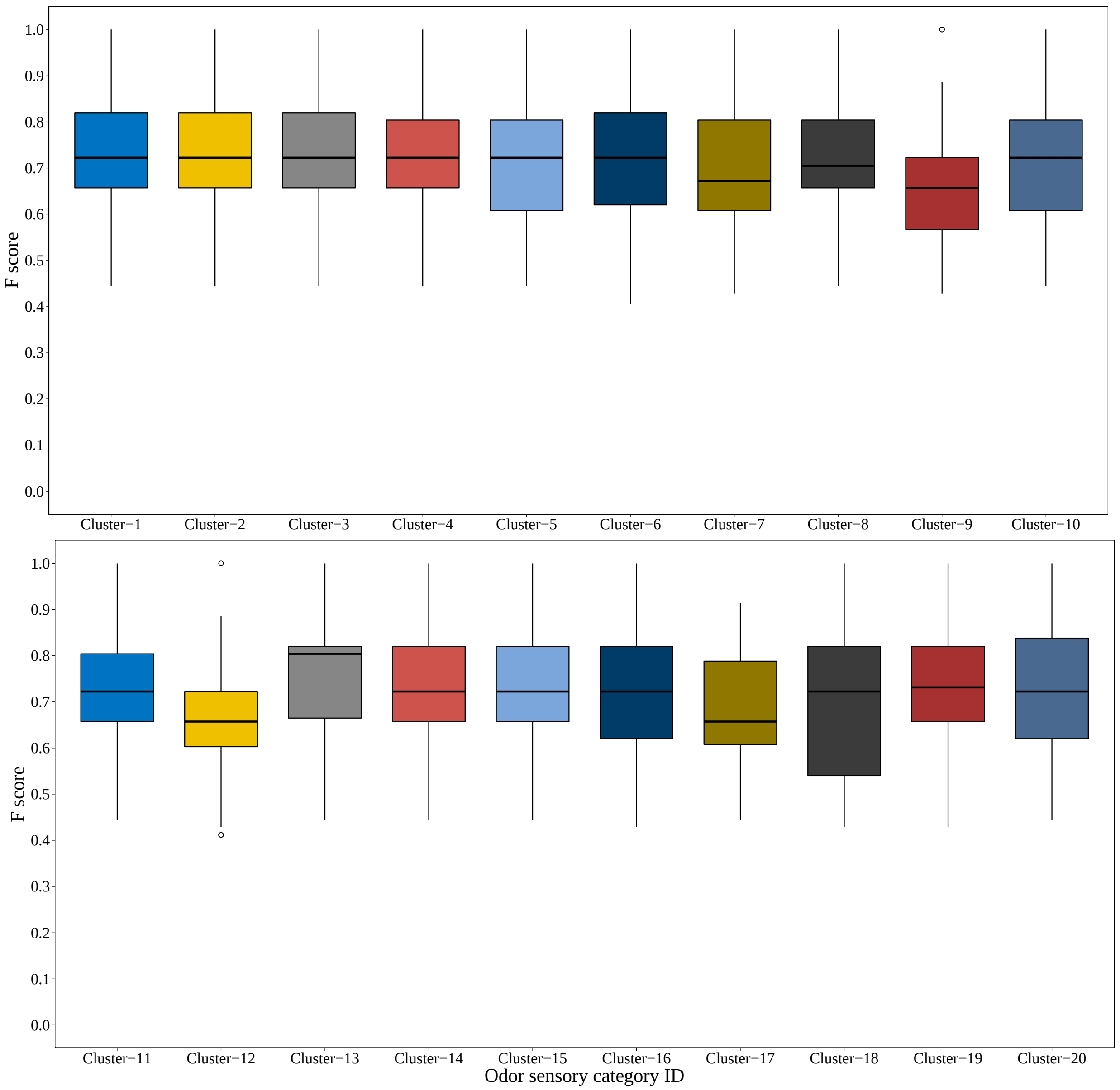

(d) F-score for MG-CNN models.

Figure S8: Comparison of odor clusters identification for area under the ROC (**a**), precision (**b**), recall (**c**) and F-score (**d**) by MG-CNN models.

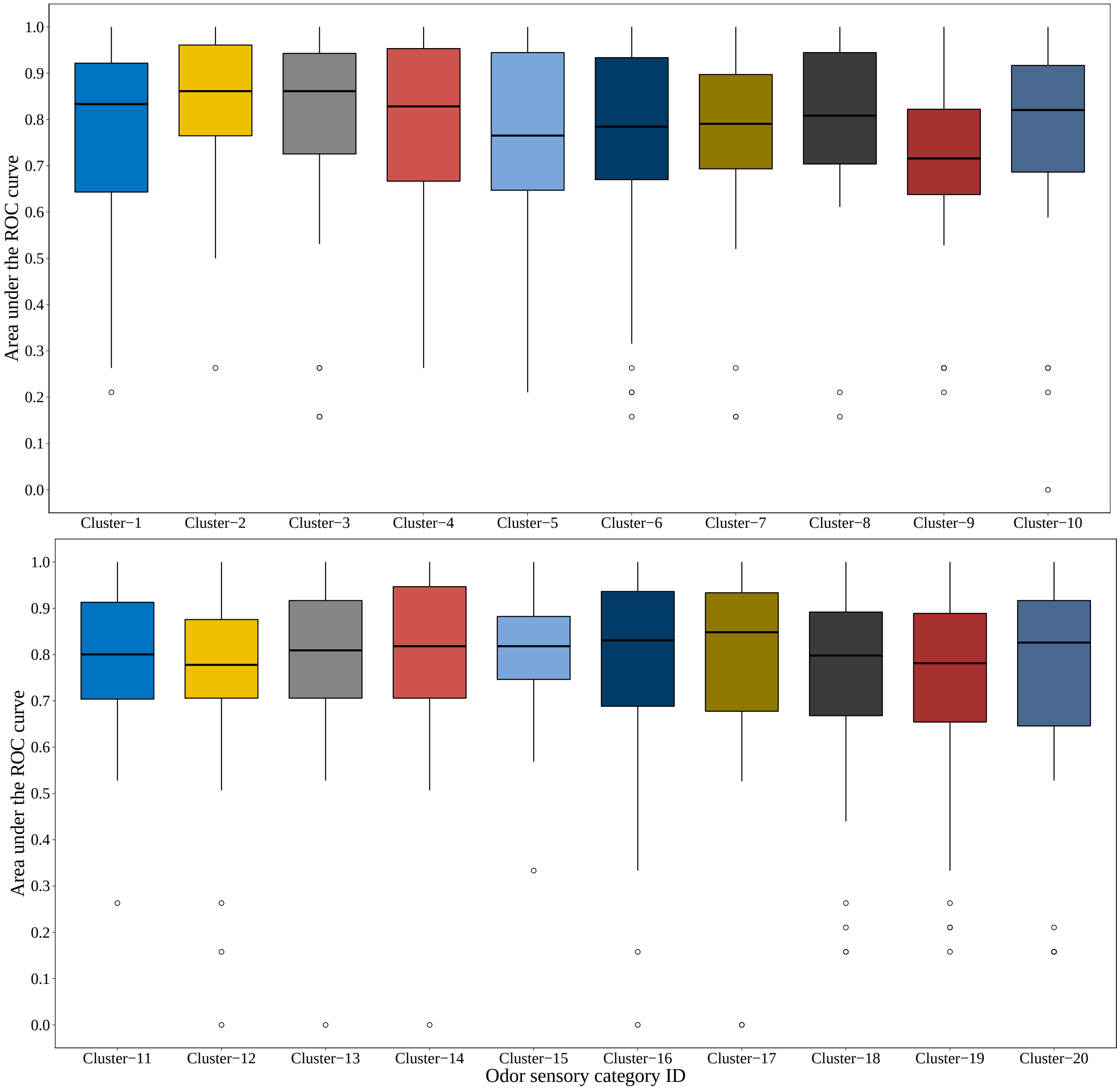

(a) Area under the ROC curve for AINN models.

Figure S9: Comparison of odor clusters identification for area under the ROC (**a**), precision (**b**), recall (**c**) and F-score (**d**) by AINN models.

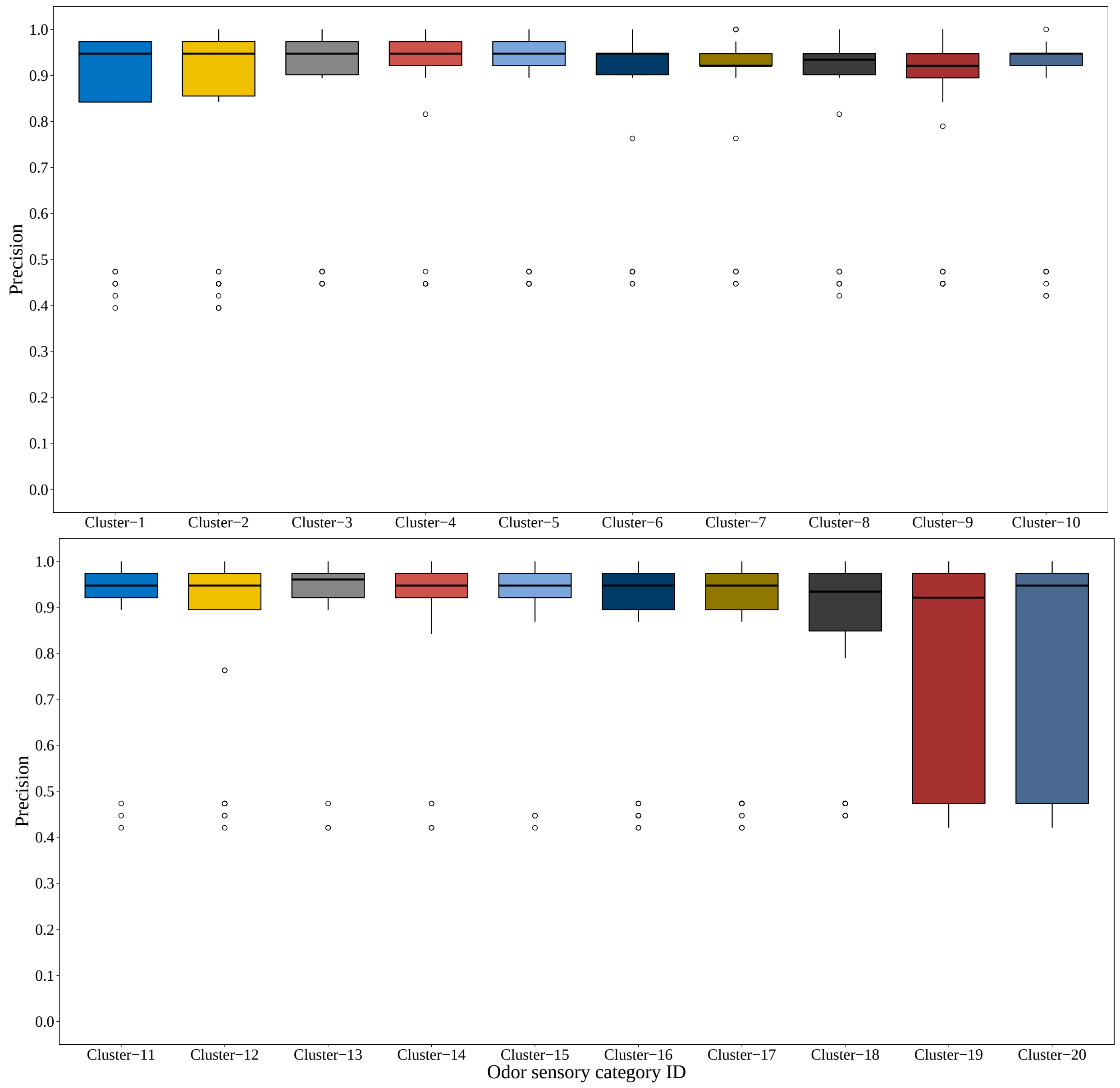

(b) Precision for AINN models.

Figure S9: Comparison of odor clusters identification for area under the ROC (**a**), precision (**b**), recall (**c**) and F-score (**d**) by AINN models.

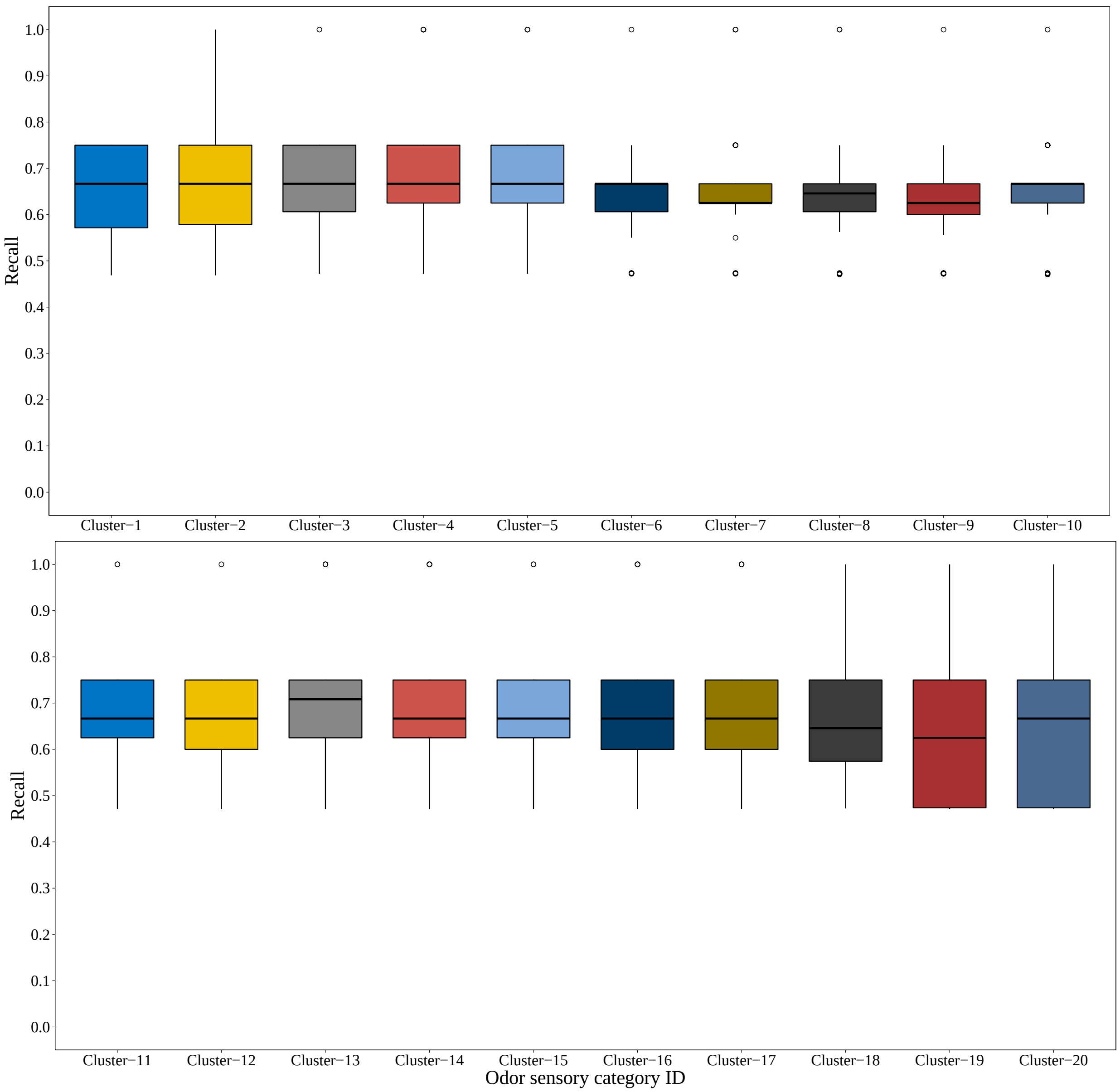

(c) Recall for AINN models.

Figure S9: Comparison of odor clusters identification for area under the ROC (**a**), precision (**b**), recall (**c**) and F-score (**d**) by AINN models.

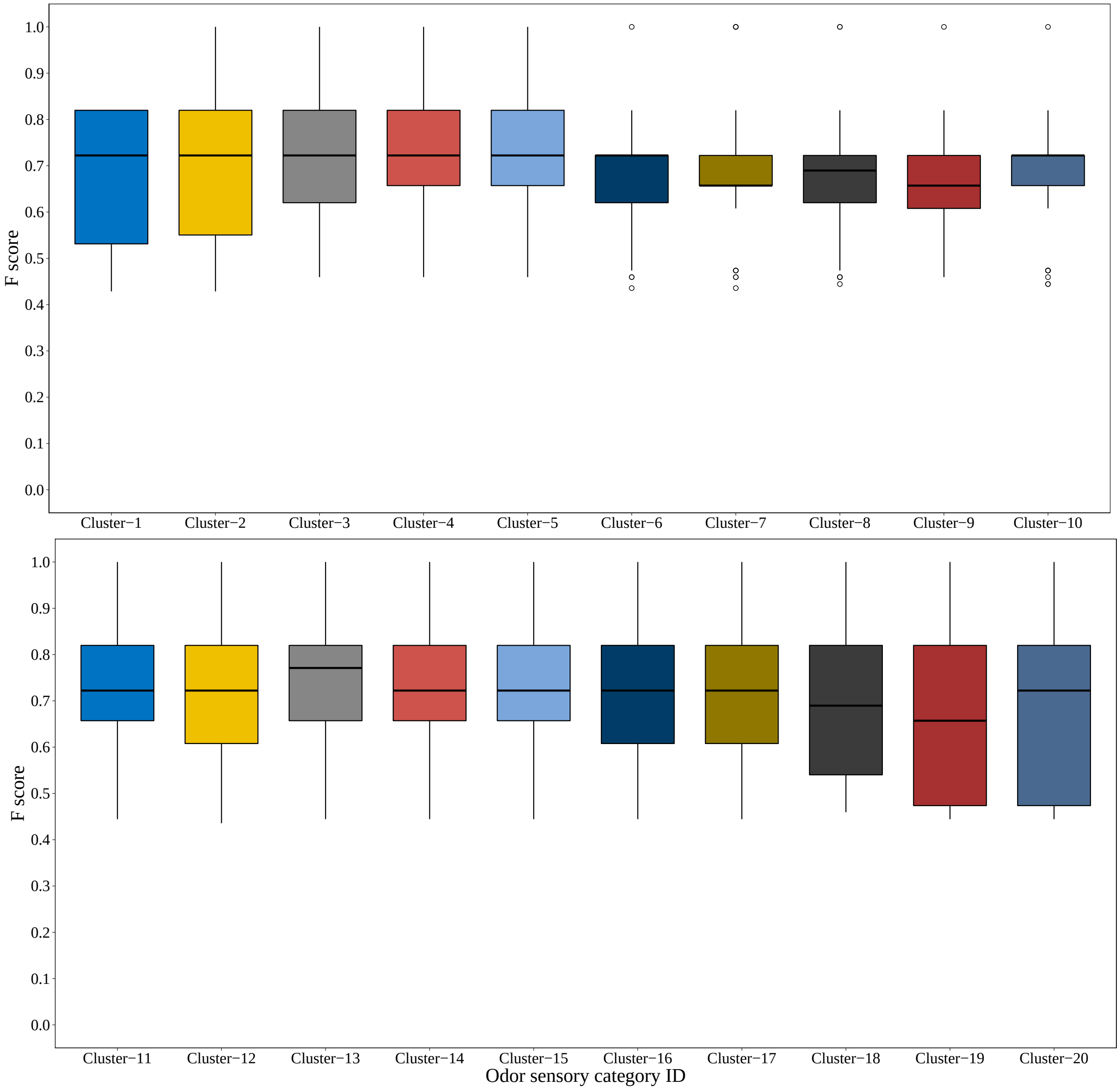

(d) F-score for AINN models.

Figure S9: Comparison of odor clusters identification for area under the ROC (**a**), precision (**b**), recall (**c**) and F-score (**d**) by AINN models.

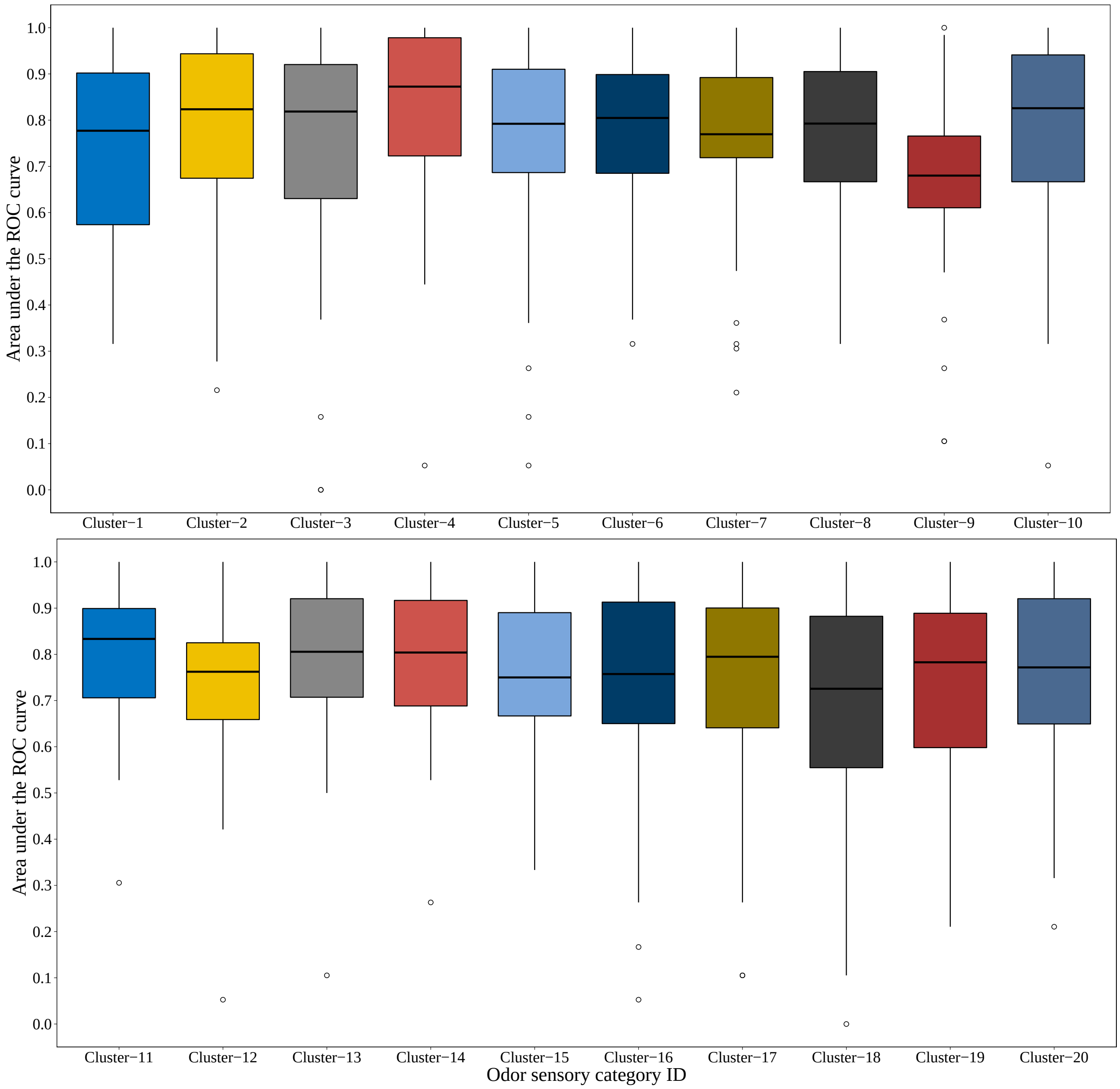

(a) Area under the ROC curve for MGTNN models.

Figure S10: Comparison of odor clusters identification for area under the ROC (**a**), precision (**b**), recall (**c**) and F-score (**d**) by MGTNN models.

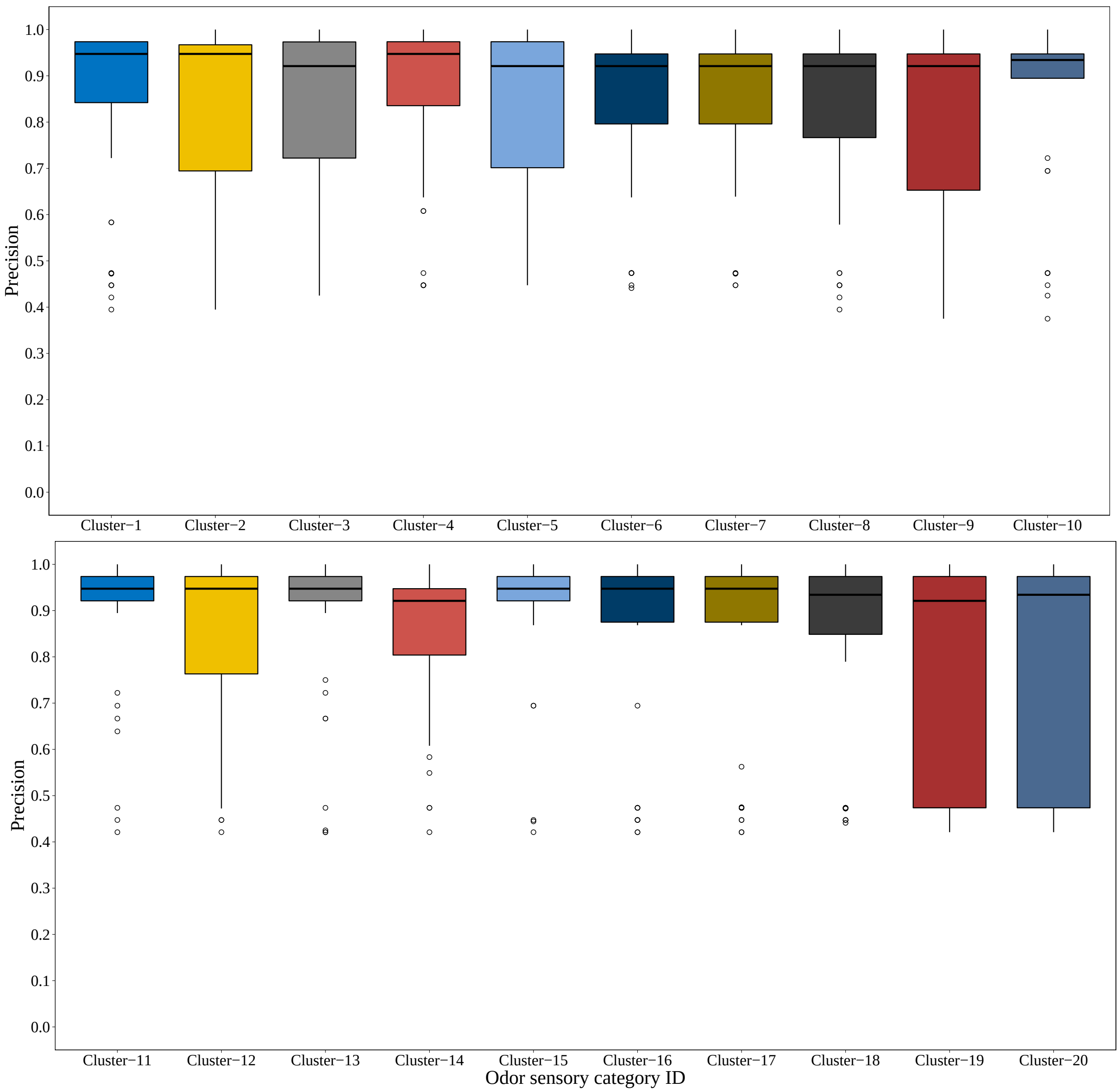

(b) Precision for MGTNN models.

Figure S10: Comparison of odor clusters identification for area under the ROC (**a**), precision (**b**), recall (**c**) and F-score (**d**) by MGTNN models.

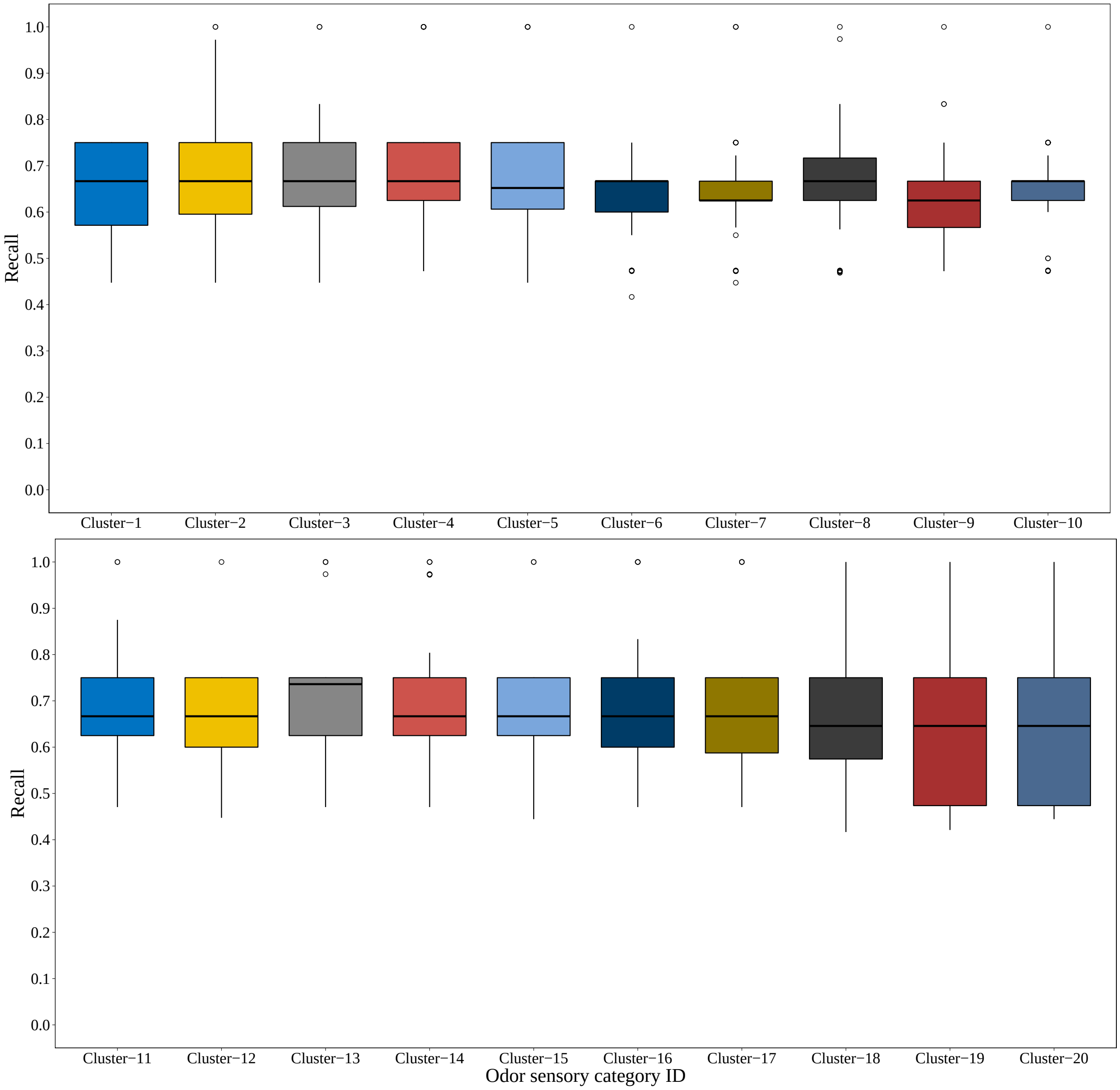

(c) Recall for MGTNN models.

Figure S10: Comparison of odor clusters identification for area under the ROC (**a**), precision (**b**), recall (**c**) and F-score (**d**) by MGTNN models.

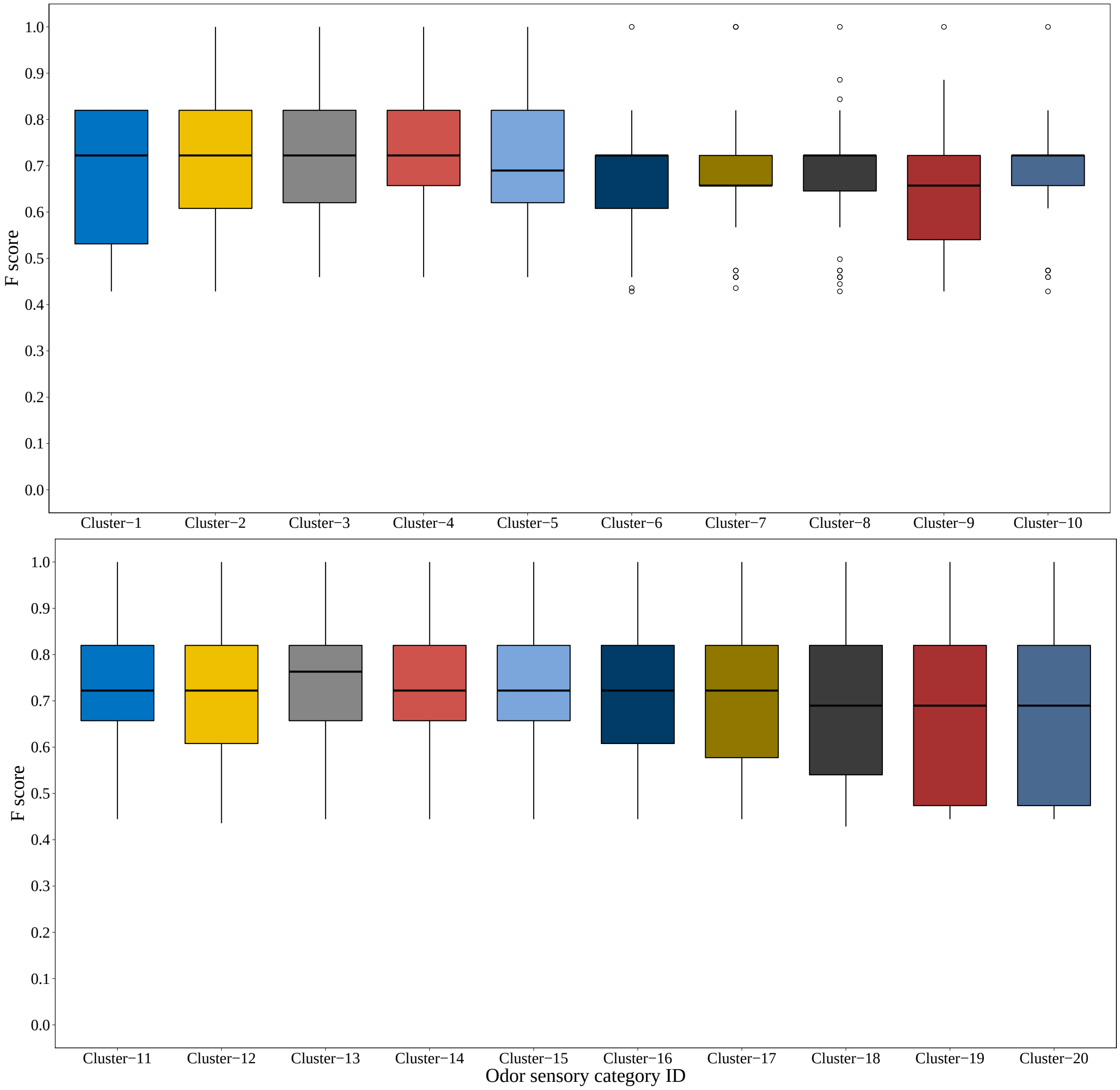

(d) F-score for MGTNN models.

Figure S10: Comparison of odor clusters identification for area under the ROC (**a**), precision (**b**), recall (**c**) and F-score (**d**) by MGTNN models.

#### Supplementary Tables

Table S1: Odor sensory categories.

| Cluster ID | Odor descriptors |
|---|---|
| Cluster-1 | butter; vanilla; powdery; buttery; tobacco; cheese; sour; rancid; coumarin; creamy; coconut; tonka; dairy; lactonic; hay; sweaty; milky; cheesy; caramellic |
| Cluster-2 | caramel; maple; sugar; fenugreek; bready |
| Cluster-3 | fruity; sweet; balsam; floral; rose; spicy; woody; green; herbal; honey |
| Cluster-4 | nutty; meaty; sulfurous; coffee; burnt; roasted; cooked; meat; chicken; savory; lamb |
| Cluster-5 | hawthorn; anisic; cherry; almond; bitter; anise; lilac; licorice; sassafrass; fennel; wintergreen; mimosa |
| Cluster-6 | balsamic; mint; aromatic; cinnamyl; pine; camphor; incense; resinous; medical; terpenic; fir; needle; labdanum; thujone |
| Cluster-7 | fatty; citrus; clean; waxy; violet; cucumber; orris; leaf; melon; watery; fresh; rind; aldehydic; soapy; ozone; marine |
| Cluster-8 | faint; juicy; winey; powerful; very; mild; bland; cognac; rum; brown; ether; estery; oil; chemical; with; odorless; alcohol; fusel; ripe; alcoholic; slightly; odor |
| Cluster-9 | plastic; strong; sharp; grassy; narcissus; cortex; leafy; musty; earthy; hyacinth; spice; foliage; mushroom; geranium; vegetable; red; metallic; orchid; weedy; galbanum; bean; cumin; mossy; pepper |
| Cluster-10 | berry; ethereal; wine; apricot; plum; banana; peach; pineapple; apple; tropical; pear; raspberry; wine-like; strawberry; grape; brandy; jam; candy |
| Cluster-11 | grapefruit; orange; petitgrain; lemon; peel; lime; mandarin; blossom; neroli; cologne; tangerine |

Table S1: Odor sensory categories.

| Cluster ID | Odor descriptors |
|---|---|
| Cluster-12 | walnut; fermented; skin; chocolate; cocoa; peanut; potato; nut; popcorn; beef; corn; chip; baked; hazelnut; toasted; grain |
| Cluster-13 | phenolic; solvent; herbaceous; minty; camphoreous; rosemary; medicinal; camphoraceous; smoky; cool; celery; caraway; mentholic; spearmint; peppery; terpene; eucalyptus; peppermint; cooling |
| Cluster-14 | alliaceous; onion; garlic; horseradish; sulfury; tomato; chrysanthemum; pungent; cabbage; mustard; radish |
| Cluster-15 | oily; rhubarb; natural; jasmin; lily; jasmine; valley; ylang; petal; ambrette; flower; gardenia; magnolia; muguet; tuberose; orangeflower; seed |
| Cluster-16 | dry; musk; civet; animal; cedar; sandalwood; amber; greasy; vetiver; dusty; ambergris; leather; wood; patchouli; acetate; old |
| Cluster-17 | warm; cinnamon; cassia; deep; tea; parsley; clove; carnation; chamomile; lavender; rooty; bergamot; sage; root; basil; bois; clary; bay; blueberry; ginger; thyme |
| Cluster-18 | fishy; ammoniacal; amine; ammonia-like; ammonia; pyridine; shrimp; seafood; pyridene |
| Cluster-19 | black; currant; mango; passion; buchu; catty |
| Cluster-20 | antiseptic; alkane; metal; almost; thiamine; rubber-like; popcorn-cracker; celery; jasminic; cardboard; vanillin; deertongue; safrole; spinach; pennyroyal; urine; woody-lactone; marigold; marjoram; calamus; turmeric; benzoin; zedoary; pomegranate; frankincense; cornmint; jonquil; sarsaparilla; kumquat; curry; chive; origanum |

Table S2: Hyper parameters tuning for MG-CNN based models

| Model name | Hidden layers | AUC-ROC | Precision | Recall | F-score |
|---|---|---|---|---|---|
| AlexNet | 2 | 0.812±0.035 | 0.862±0.04 | 0.661±0.029 | 0.695±0.031 |
| AlexNet | 3 | 0.811±0.033 | 0.862±0.034 | 0.66±0.027 | 0.694±0.03 |
| AlexNet | 4 | 0.812±0.035 | 0.859±0.04 | 0.658±0.028 | 0.692±0.032 |
| AlexNet | 5 | 0.808±0.036 | 0.866±0.034 | 0.659±0.027 | 0.695±0.03 |
| AlexNet | 6 | 0.81±0.034 | 0.86±0.037 | 0.66±0.027 | 0.694±0.03 |
| Pre-trained AlexNet | 2 | 0.86±0.031 | 0.824±0.044 | 0.703±0.04 | 0.721±0.036 |
| Pre-trained AlexNet | 3 | 0.869±0.033 | 0.83±0.041 | 0.704±0.037 | 0.722±0.035 |
| Pre-trained AlexNet | 4 | 0.871±0.028 | 0.834±0.042 | 0.706±0.035 | 0.726±0.033 |
| Pre-trained AlexNet | 5 | 0.873±0.029 | 0.822±0.042 | 0.707±0.035 | 0.723±0.033 |
| Pre-trained AlexNet | 6 | 0.871±0.031 | 0.83±0.043 | 0.704±0.037 | 0.723±0.034 |
| VGG | 2 | 0.815±0.031 | 0.912±0.019 | 0.667±0.017 | 0.715±0.02 |
| VGG | 3 | 0.829±0.028 | 0.899±0.019 | 0.665±0.017 | 0.713±0.02 |
| VGG | 4 | 0.82±0.021 | 0.912±0.019 | 0.667±0.017 | 0.715±0.02 |
| VGG | 5 | 0.818±0.014 | 0.912±0.019 | 0.667±0.017 | 0.715±0.02 |
| VGG | 6 | 0.819±0.015 | 0.912±0.019 | 0.667±0.017 | 0.715±0.02 |
| Pre-trained VGG | 2 | 0.857±0.034 | 0.809±0.046 | 0.697±0.032 | 0.712±0.03 |
| Pre-trained VGG | 3 | 0.869±0.029 | 0.815±0.047 | 0.702±0.039 | 0.716±0.037 |
| Pre-trained VGG | 4 | 0.87±0.03 | 0.822±0.04 | 0.699±0.036 | 0.717±0.032 |
| Pre-trained VGG | 5 | 0.872±0.028 | 0.824±0.043 | 0.7±0.035 | 0.719±0.034 |
| Pre-trained VGG | 6 | 0.875±0.028 | 0.827±0.036 | 0.698±0.036 | 0.716±0.033 |
| DenseNet | 2 | 0.775±0.029 | 0.856±0.043 | 0.676±0.025 | 0.715±0.032 |
| DenseNet | 3 | 0.81±0.013 | 0.906±0.011 | 0.669±0.021 | 0.717±0.021 |
| DenseNet | 4 | 0.783±0.033 | 0.852±0.103 | 0.664±0.02 | 0.703±0.037 |
| DenseNet | 5 | 0.823±0.013 | 0.906±0.028 | 0.666±0.018 | 0.714±0.022 |
| DenseNet | 6 | 0.833±0.02 | 0.891±0.02 | 0.669±0.018 | 0.715±0.021 |

Table S2: Hyper parameters tuning for MG-CNN based models

| Model name | Hidden layers | AUC-ROC | Precision | Recall | F-score |
|---|---|---|---|---|---|
| Pre-trained DenseNet | 2 | 0.873±0.031 | 0.829±0.04 | 0.701±0.036 | 0.72±0.035 |
| Pre-trained DenseNet | 3 | 0.872±0.027 | 0.829±0.036 | 0.702±0.035 | 0.72±0.034 |
| Pre-trained DenseNet | 4 | 0.877±0.029 | 0.823±0.037 | 0.697±0.036 | 0.716±0.035 |
| Pre-trained DenseNet | 5 | 0.872±0.028 | 0.815±0.037 | 0.7±0.036 | 0.715±0.034 |
| Pre-trained DenseNet | 6 | 0.874±0.027 | 0.818±0.037 | 0.708±0.033 | 0.724±0.031 |
| ResNet | 2 | 0.753±0.037 | 0.711±0.038 | 0.657±0.033 | 0.657±0.031 |
| ResNet | 3 | 0.779±0.032 | 0.745±0.049 | 0.665±0.03 | 0.672±0.032 |
| ResNet | 4 | 0.774±0.035 | 0.78±0.045 | 0.655±0.026 | 0.672±0.027 |
| ResNet | 5 | 0.788±0.03 | 0.802±0.055 | 0.658±0.031 | 0.68±0.033 |
| ResNet | 6 | 0.792±0.037 | 0.803±0.056 | 0.662±0.03 | 0.684±0.032 |
| Pre-trained ResNet | 2 | 0.876±0.024 | 0.842±0.034 | 0.696±0.034 | 0.719±0.032 |
| Pre-trained ResNet | 3 | 0.876±0.027 | 0.837±0.043 | 0.698±0.039 | 0.719±0.039 |
| Pre-trained ResNet | 4 | 0.876±0.027 | 0.821±0.038 | 0.703±0.03 | 0.72±0.03 |
| Pre-trained ResNet | 5 | 0.876±0.029 | 0.823±0.045 | 0.703±0.036 | 0.72±0.037 |
| Pre-trained ResNet | 6 | 0.877±0.028 | 0.822±0.037 | 0.71±0.028 | 0.726±0.028 |

Table S3: Hyper parameters tuning for MGTNN based models.

| Model name | Hidden layers | AUC-ROC | Precision | Recall | F-score |
|---|---|---|---|---|---|
| Pre-trained GCN atom | 2 | 0.805±0.036 | 0.845±0.046 | 0.661±0.027 | 0.692±0.03 |
| Pre-trained GCN atom | 3 | 0.812±0.03 | 0.852±0.033 | 0.66±0.027 | 0.693±0.03 |
| Pre-trained GCN atom | 4 | 0.803±0.035 | 0.859±0.035 | 0.662±0.028 | 0.696±0.032 |
| Pre-trained GCN atom | 5 | 0.801±0.035 | 0.852±0.038 | 0.666±0.027 | 0.697±0.03 |
| Pre-trained GCN atom | 6 | 0.809±0.033 | 0.855±0.037 | 0.661±0.027 | 0.693±0.029 |
| Pre-trained GCN atom+bond | 2 | 0.804±0.034 | 0.845±0.037 | 0.659±0.028 | 0.691±0.031 |
| Pre-trained GCN atom+bond | 3 | 0.807±0.033 | 0.846±0.034 | 0.659±0.027 | 0.69±0.029 |
| Pre-trained GCN atom+bond | 4 | 0.807±0.033 | 0.853±0.037 | 0.659±0.027 | 0.691±0.03 |
| Pre-trained GCN atom+bond | 5 | 0.808±0.036 | 0.849±0.035 | 0.659±0.026 | 0.691±0.029 |
| Pre-trained GCN atom+bond | 6 | 0.804±0.036 | 0.855±0.036 | 0.658±0.025 | 0.692±0.029 |
| Pre-trained GCN atom+bond | 7 | 0.813±0.035 | 0.856±0.037 | 0.663±0.028 | 0.696±0.032 |
| Pre-trained GCN atom+bond | 8 | 0.809±0.036 | 0.85±0.042 | 0.661±0.029 | 0.693±0.033 |
| Pre-trained GCN bond | 2 | 0.804±0.035 | 0.838±0.046 | 0.659±0.027 | 0.689±0.031 |
| Pre-trained GCN bond | 3 | 0.81±0.03 | 0.85±0.039 | 0.661±0.029 | 0.693±0.033 |
| Pre-trained GCN bond | 4 | 0.805±0.034 | 0.856±0.032 | 0.661±0.026 | 0.694±0.028 |
| Pre-trained GCN bond | 5 | 0.806±0.037 | 0.857±0.036 | 0.663±0.027 | 0.696±0.03 |
| Pre-trained GCN bond | 6 | 0.81±0.034 | 0.852±0.037 | 0.661±0.028 | 0.693±0.031 |

Table S4: Hyper parameters tuning for AINN based models.

| Model name | Embedded dim | AUC-ROC | Precision | Recall | F-score |
|---|---|---|---|---|---|
| AINN only (2D) | 32 | 0.785±0.039 | 0.864±0.042 | 0.661±0.026 | 0.696±0.030 |
| AINN only (2D) | 64 | 0.786±0.039 | 0.867±0.034 | 0.660±0.027 | 0.695±0.031 |
| AINN only (2D) | 128 | 0.788±0.040 | 0.867±0.034 | 0.660±0.027 | 0.696±0.031 |
| AINN only (2D) | 256 | 0.792±0.039 | 0.867±0.034 | 0.660±0.027 | 0.695±0.031 |
| AINN only (2D) | 512 | 0.790±0.041 | 0.867±0.034 | 0.660±0.027 | 0.695±0.031 |
| AINN only (2D) | 32 | 0.786±0.045 | 0.865±0.037 | 0.656±0.028 | 0.692±0.032 |
| AINN only (2D) | 64 | 0.781±0.046 | 0.865±0.037 | 0.656±0.028 | 0.692±0.032 |
| AINN only (2D) | 128 | 0.786±0.045 | 0.865±0.037 | 0.656±0.028 | 0.692±0.032 |
| AINN only (2D) | 256 | 0.785±0.044 | 0.865±0.037 | 0.656±0.028 | 0.692±0.032 |
| AINN only (2D) | 512 | 0.787±0.045 | 0.864±0.037 | 0.656±0.028 | 0.692±0.032 |
| AINN (2D) | 32 | 0.803±0.038 | 0.863±0.037 | 0.659±0.027 | 0.694±0.031 |
| AINN (2D) | 64 | 0.794±0.037 | 0.864±0.037 | 0.659±0.027 | 0.695±0.031 |
| AINN (2D) | 128 | 0.805±0.038 | 0.863±0.036 | 0.659±0.027 | 0.694±0.03 |
| AINN (2D) | 256 | 0.802±0.038 | 0.860±0.040 | 0.662±0.026 | 0.696±0.03 |
| AINN (2D) | 512 | 0.807±0.035 | 0.861±0.038 | 0.662±0.027 | 0.696±0.03 |
| AINN (3D) | 32 | 0.797±0.035 | 0.838±0.035 | 0.656±0.027 | 0.683±0.029 |
| AINN (3D) | 64 | 0.782±0.043 | 0.832±0.045 | 0.656±0.028 | 0.685±0.03 |
| AINN (3D) | 128 | 0.794±0.037 | 0.852±0.05 | 0.657±0.028 | 0.689±0.033 |
| AINN (3D) | 256 | 0.800±0.036 | 0.832±0.046 | 0.655±0.030 | 0.683±0.034 |
| AINN (3D) | 512 | 0.781±0.037 | 0.817±0.044 | 0.660±0.035 | 0.675±0.033 |
| ResAINN only (2D) | 32 | 0.793±0.040 | 0.867±0.034 | 0.660±0.027 | 0.695±0.031 |
| ResAINN only (2D) | 64 | 0.797±0.038 | 0.867±0.034 | 0.660±0.027 | 0.695±0.031 |
| ResAINN only (2D) | 128 | 0.798±0.038 | 0.867±0.034 | 0.660±0.027 | 0.695±0.030 |
| ResAINN only (2D) | 256 | 0.797±0.039 | 0.866±0.034 | 0.659±0.027 | 0.695±0.030 |
| ResAINN only (2D) | 512 | 0.797±0.038 | 0.867±0.034 | 0.659±0.027 | 0.695±0.031 |

Table S4: Hyper parameters tuning for AINN based models.

| Model name | Embedded dim | AUC-ROC | Precision | Recall | F-score |
|---|---|---|---|---|---|
| ResAINN only (3D) | 32 | 0.791±0.044 | 0.865±0.037 | 0.656±0.028 | 0.692±0.032 |
| ResAINN only (3D) | 64 | 0.793±0.043 | 0.865±0.037 | 0.656±0.028 | 0.692±0.032 |
| ResAINN only (3D) | 128 | 0.795±0.041 | 0.866±0.037 | 0.658±0.028 | 0.693±0.032 |
| ResAINN only (3D) | 256 | 0.796±0.042 | 0.865±0.037 | 0.658±0.028 | 0.693±0.032 |
| ResAINN only (3D) | 512 | 0.796±0.041 | 0.860±0.039 | 0.657±0.026 | 0.690±0.030 |
| ResAINN (2D) | 32 | 0.802±0.036 | 0.863±0.035 | 0.660±0.027 | 0.695±0.030 |
| ResAINN (2D) | 64 | 0.795±0.038 | 0.865±0.036 | 0.660±0.027 | 0.695±0.031 |
| ResAINN (2D) | 128 | 0.805±0.037 | 0.864±0.034 | 0.659±0.027 | 0.695±0.030 |
| ResAINN (2D) | 256 | 0.801±0.038 | 0.863±0.037 | 0.660±0.028 | 0.695±0.032 |
| ResAINN (2D) | 512 | 0.806±0.035 | 0.859±0.039 | 0.661±0.026 | 0.694±0.030 |
| ResAINN (3D) | 32 | 0.794±0.035 | 0.831±0.053 | 0.659±0.028 | 0.684±0.032 |
| ResAINN (3D) | 64 | 0.783±0.042 | 0.840±0.042 | 0.655±0.029 | 0.686±0.032 |
| ResAINN (3D) | 128 | 0.794±0.036 | 0.856±0.050 | 0.656±0.029 | 0.690±0.036 |
| ResAINN (3D) | 256 | 0.797±0.034 | 0.840±0.041 | 0.663±0.032 | 0.689±0.034 |
| ResAINN (3D) | 512 | 0.783±0.038 | 0.813±0.045 | 0.661±0.036 | 0.674±0.034 |
